# Design and optimization of a kinase-controlled allosteric switch

**DOI:** 10.1101/2025.04.15.648940

**Authors:** Qinhao Cao, Jared E. Toettcher

## Abstract

Post-translational control enables rapid and precise regulation of cell behavior. Despite these advantages, general strategies to build phosphorylation-based synthetic circuits are limited. Here, we reasoned that engineered allostery, a technique that has been applied to design light- and chemically-gated protein switches, could also be used to engineer phosphorylation-controlled protein switches (phospho-switches). Using an allosterically controllable Gal4 transcription factor as a scaffold, we show that a classic kinase FRET biosensor architecture can be used as a starting point for phospho-switch design. We optimize all features of the phospho-switch to develop an ERK-controlled transcription factor with a 20-fold phosphorylation-dependent change in transcriptional output. Our switch outperforms the *c-fos* promoter, a classic ERK-responsive transcriptional biosensor, in selectivity and sensitivity. We further show that our switch architecture can be generalized to other input kinases and allosterically controlled targets. This work provides a general platform for a new generation of kinase-responsive tools for biosensing and synthetic biology applications.

## Introduction

Cells rely on protein phosphorylation as a primary carrier of intracellular information. Protein kinases are activated by a variety of stimuli and regulate diverse processes including cytoskeletal assembly, metabolism, protein localization, and gene expression. Phosphorylation can also link multiple kinases and substrates into networks that can perform diverse signal processing capabilities, including signal integration^1^, amplification^2^, and feedback control^3^. Yet while bioengineers have long been able to build synthetic transcriptional circuits^4,5^, our ability to engineer post-translational networks has been far more limited.

Recent studies have already begun to demonstrate the power of synthetic post-translational control for sensing and rewiring oncogenic states, constructing neural networks to perform information processing in cells, and improving engineered cell-based therapies^6–9^. Yet each of these studies relied on engineered protease-substrate interactions to perform stimulus-dependent irreversible cleavage of an effector domain^10,11^. In contrast to protein cleavage, the reversibility of protein phosphorylation presents an opportunity to provide bidirectional control for more fine-tuned responses in space and time. Moreover, synthetic phosphorylation-based logic would make it possible to directly interface with the kinome to sense kinase-encoded cell states and trigger desired responses.

Engineering proteins whose activity is controlled by defined phosphorylation events is challenging, but some studies have begun to address this challenge. A pioneering study demonstrated that a novel phosphorylation site could be introduced into the *Saccharomyces cerevisiae* kinase Kss1 to regulate its activity^12^. Phosphorylation-dependent binding has also been used in *S. cerevisiae* to re-localize cytosolic proteins to the plasma membrane, activating cellular processes in cases where membrane localization is essential for function^13^. Most recently, synthetic phosphorylation relays were constructed in mammalian cells using combinations of engineered tyrosine kinases, phospho-tyrosine (pTyr) peptides, and SH2 pTyr-binding domains^14^. General methods for phosphorylation-dependent control of protein function could power advances in synthetic biology by enabling the production of compact phosphorylation-based intracellular circuits.

Engineered allostery is a powerful concept for achieving stimulus-dependent control over protein function. In engineered allostery, a stimulus-regulated protein domain (switch) is inserted at a particular site in a target protein, such that a conformational change in the switch alters the target protein’s function (**Fig. 1A**). Several domains, including the light-sensitive AsLOV2 domain and ligand-responsive uniRapR domain, can act as suitable stimulus-dependent switches, changing conformation in response to illumination or addition of a small molecule (**Fig. 1B**)^15,16^. However, no switches are yet available to respond to protein phosphorylation. A kinase-controlled phosphorylation-dependent switch could immediately confer phospho-regulation to the diverse set of target proteins previously engineered to be controlled by light or small molecule addition.

**Figure 1.**
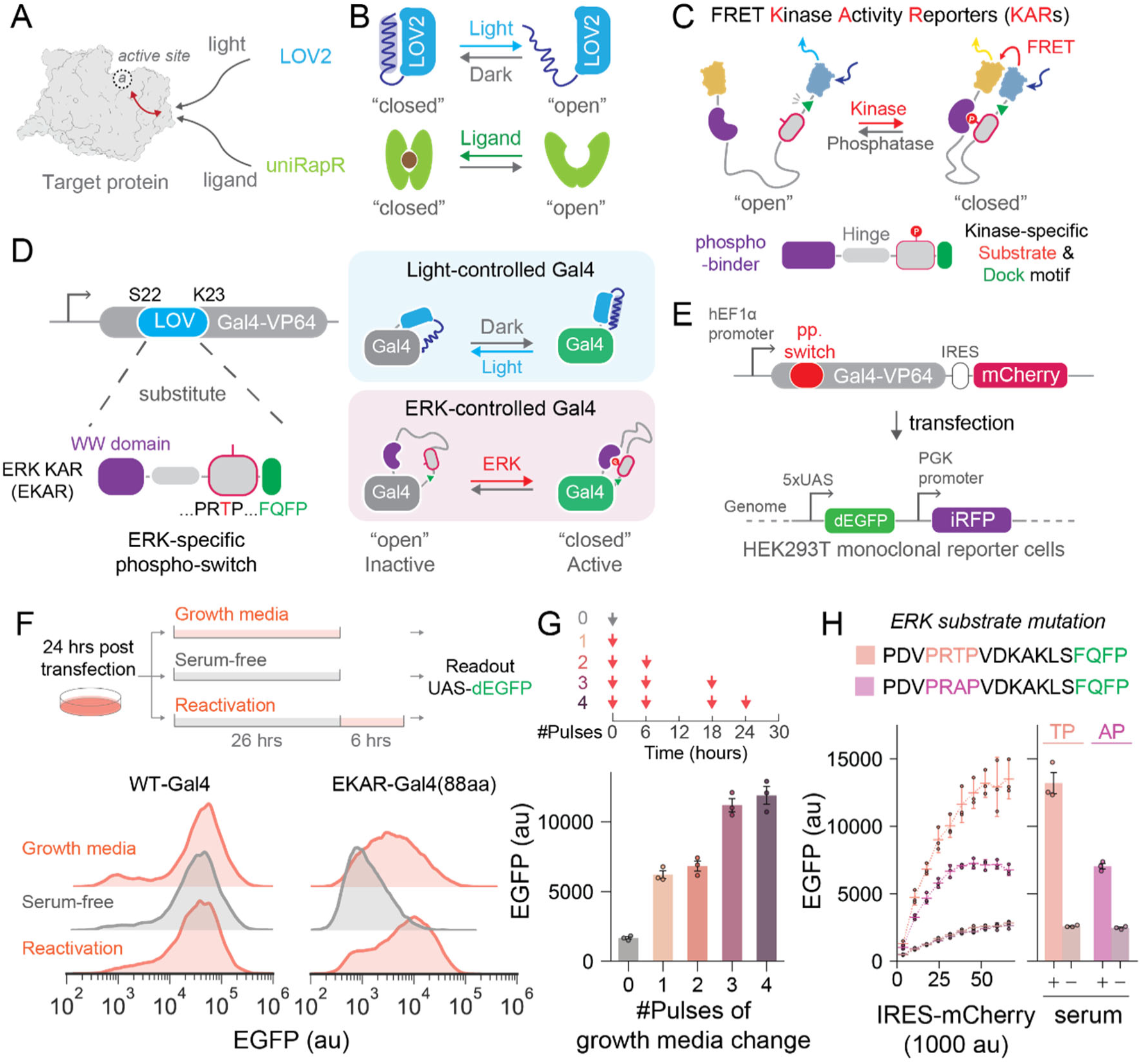
Repurposing FRET kinase activity biosensors as allosteric switches. (**A**) Schematic of engineered allosteric control of protein function. A stimulus-regulated protein domain, like light-sensitive AsLOV2 (blue) or ligand-sensitive uniRapR (green), is inserted into an allosteric site in a target protein and coupled to its function. (**B**) Mechanism of action for existing light/ligand-sensitive switches, which undergo a stimulus-dependent conformational change. (**C**) Schematic of FRET-based Kinase Activity Reporters (KARs). Fluorescent proteins on the N and C termini are linked by a phospho-binding domain (purple), a hinge region (gray) and a kinase-specific substrate (red) with docking motif (green). (**D**) The Gal4 DNA binding domain was previously made light-switchable by AsLOV2 insertion between Ser22 and Lys23. Replacing AsLOV2 with the ERK KAR (EKAR) produces a candidate ERK-controlled phosphoGal4 (EKAR-Gal4). (**E**) Overview of EKAR-Gal4 testing pipeline. EKAR-Gal4 was transiently expressed with an expression marker (IRES-mCherry) in a monoclonal 5xUAS-dEGFP HEK293T reporter cell line and EGFP expression was monitored in ERK-ON and ERK-OFF conditions. (**F**) Serum-dependent GFP expression of reporter cells transfected with Gal4-VP64 (WT-Gal4) or EKAR-Gal4. Cells were incubated in serum-free or growth media for 26 h before measurement. Reactivation was measured 6 h after switching back to growth media. Histogram represents >10000 cells combined from 3 replicates. (**G**) GFP expression in response to repeated growth media stimulation in EKAR-Gal4 expressing reporter cells. Error bars represent mean ± s.e.m (n = 3 replicates). (**H**) GFP expression in reporter cells expressing EKAR-Gal4 (PRTP) or its non-phosphorylatable T-to-A mutant (PRAP) in serum-free or growth media conditions. Curves present GFP expression across a range of TF expression levels (4 curves shown, with the serum-free conditions overlapping for PRTP and PRAP variants). Error bars on the curve represent the mean ± 95% confidence interval. Error bars in the right bar graph represent mean ± s.e.m (n = 3 replicates).

Here, we discover that a commonly used Förster resonance energy transfer (FRET) kinase biosensor provides a starting point for the development of a kinase-controlled allosteric switch. We show that the EKAR biosensor^17,18^ for the Extracellular Signal-Related Kinase (ERK), can function as a prototype kinase-controlled switch when inserted into a Gal4 transcription factor at a previously identified allosteric site. We systematically optimize each component of the switch (hinge, phospho-peptide, and phospho-binding domain) to substantially improve its potency and dynamic range, revealing design principles for achieving phosphorylation-dependent allostery that differ from FRET biosensors. Our final switch design results in a >20-fold change in transcriptional output between phosphorylated and unphosphorylated states. We show that our synthetic kinase-controlled transcription factor functions as an excellent biosensor of ERK kinase activity, outperforming the FOS promoter in sensitivity and selectivity for ERK over other stimuli. Our switch generalizes to multiple kinases and allosterically-controlled target proteins, including a JNK-controlled transcription factor and an ERK-controlled actin nanobody. Overall, this strategy expands the synthetic biologist’s toolbox of post-translationally regulated processes to a diverse suite of kinases and allosterically-controlled target proteins with potential applications in biosensor design, drug screening, and cell-based therapy.

## Results

### Repurposing FRET-based kinase activity reporters as allosteric switches

Our strategy for achieving kinase-triggered allosteric control requires a phosphorylation-regulated sensory domain (phospho-switch) that can regulate the conformation of a target protein based on the activity of a specific upstream kinase. An ideal phospho-switch would satisfy a few criteria. First, phosphorylation should trigger an open-to-closed conformational change that alters the distance between the switch’s N and C termini, in a similar manner to successful chemogenetic and optogenetic allosteric switches^15,16^. Second, it should be rapidly and reversibly regulated by phosphorylation from a defined kinase of interest. Third, the switch design should be modular and generalizable to multiple kinase inputs.

We reasoned that FRET-based kinase activity reporters (KARs) provide an attractive starting point for satisfying each of these criteria. The sensory region in KARs consists of a phosphorylatable substrate, a flexible hinge, and a phosphate-binding domain (phospho-binder)^19^. Binding between the phosphorylated substrate and phospho-binder brings fluorescent proteins on the KAR’s N and C termini into proximity to produce a change in fluorescence (**Fig. 1C**). KARs have been shown to rapidly and reversibly report on kinase activity, and due to their modular design, many KARs targeting diverse kinases (e.g., ERK, PKA, SRC) have already been developed and validated^20^. We thus hypothesized that KAR sensory domains could be repurposed as prototype phospho-switches to allosterically control target proteins of interest. We focused on the EKAR-EV biosensor for the ERK kinase, a cell signaling protein that is activated by extracellular growth factors and whose activity stimulates cell growth and differentiation during development, tissue regeneration, and cancer progression. The EKAR-EV biosensor contains an ERK-responsive sensory domain that includes an ERK-phosphorylatable peptide (amino acids PRTP) and docking motif (FQFP), an 88 amino acid Gly/Ser/Ala hinge, and the Pin1 WW phospho-binding domain^17,18^.

We chose an engineered transcription factor formed from the Gal4 DNA binding domain and the VP64 transactivation domain as a model target protein for implementing ERK-based allosteric control. In a recent study, we developed a light-controlled Gal4-VP64 variant by inserting the photoswitchable AsLOV2 domain between positions Ser22 and Lys23 (SK22) inside the Gal4 DNA-binding domain (**Fig. 1D**). The resulting OptoGal4 system exhibits potent allosteric control, with a >100-fold change in transcription activity between blue light illumination and dark conditions^21^. We replaced the AsLOV2 module with the EKAR sensory domain to test whether the light-responsive protein could instead be made to sense a change in phosphorylation state, terming the resulting fusion protein EKAR-Gal4. When ERK is active, the WW domain would be expected to bind the phosphorylated peptide, resulting in a closed switch and a transcriptionally active Gal4 (**Fig. 1D**).

To assess the kinase-controlled activity of EKAR-Gal4, we used a previously developed clonal HEK293T cell line in which EGFP expression is driven by a Gal4-responsive *5xUAS* promoter (*5xUAS*-dEGFP) (**Fig. 1E**)^21^. We transiently transfected these cells with an EKAR-Gal4 plasmid that also included IRES-mCherry marker to assess cells’ relative Gal4 expression levels (**Fig. 1E**). We then incubated cells in either serum-free media to induce an ERK-OFF state or serum-containing growth media to maintain the ERK-ON state (see **Methods**; for flow cytometry analysis pipeline see **Fig. S1**). We observed approximately three-fold higher EGFP expression from cells incubated for 26 h in growth media compared to serum-free media (**Fig. 1F**). We also found that EKAR-Gal4-induced EGFP expression could be reactivated within 6 h after a switch back from serum-free to growth media (**Fig. 1F**). In contrast, cells expressing an unmodified Gal4-VP64 transcription factor (“WT-Gal4”) exhibited high levels of EGFP induction in both serum-free and growth media conditions (**Fig. 1F**).

ERK signaling can be dynamic, with cells only transiently entering an ERK-high state after growth factor stimulation^22–24^, which might limit the extent of Gal4 transcriptional activity in our system. To test if this was the case in HEK293Ts, we characterized ERK dynamics after growth media stimulation using an ERK biosensor (ErkKTR) that is exported from the nucleus in response to ERK activation^25^. We found that ERK is only transiently activated for ∼1 h by growth media stimulation in HEK293 cells and that after adaptation, cells can be activated again by exchanging the media for fresh growth media (**Fig. S2**). Based on these results, we hypothesized that a series of growth media exchanges might produce a higher Gal4 response. Indeed, the level of EKAR-Gal4-induced EGFP expression was correlated to the number of growth media exchanges delivered within 30 hours (**Fig. 1G**). For all subsequent experiments, we exchanged media at 0, 6, 18, 24 h prior to quantification of EKAR-Gal4 transcriptional output at 30 h.

We next sought to confirm that the change in EKAR-Gal4 transcriptional activity depends on the phospho-peptide’s phosphorylation state. We made an EKAR-Gal4 variant with a threonine-to-alanine mutation in the consensus ERK phosphorylation site (PRTP to PRAP). In ERK-ON conditions, the PRAP mutant exhibited reduced transcriptional activity compared to the PRTP variant, but higher activity than cells in serum-free media (**Fig. 1H**). This data suggests that EKAR-Gal4 transcriptional activity is controlled by serum factors beyond a single phosphorylation event, an undesirable effect for our system and a primary target for optimization in subsequent experiments.

In summary, our initial experiments suggest that EKAR insertion is a promising starting point for the development of an ERK-controlled phospho-switch. However, further optimization is essential to improve its low potency, limited dynamic range, and residual activity in the T-to-A mutant. For version control purposes, we term this original EKAR as “phospho-switch v0” and the corresponding EKAR-Gal4 as “ERK-phosphoGal4v0”.

### Hinge optimization reveals allosteric switch design principles

As a first target for phospho-switch optimization, we examined the hinge region that links the phosphorylatable peptide to the WW phospho-binding domain. Prior studies of FRET kinase biosensors revealed that varying the length of this hinge can have substantial effects on biosensor efficacy, with longer hinges typically producing a lower baseline of FRET activity and higher biosensor gain^18^. However, the optimal hinge for an allosteric switch may differ substantially from that of a FRET biosensor. We thus set out to explore a broad design space of hinge sequences, focusing on improving the maximum activation in ERK-ON conditions and dynamic range between ERK-ON and ERK-OFF conditions as design goals.

We constructed a combinatorial Golden-Gate cloning system (QCTK) for rapidly generating allosteric switch variants (**Fig. 2A**; see **Methods**). We designed level 1 “a”, “b”, and “c” parts consisting of phospho-binding domains, hinge sequences, and phosphorylatable substrates, respectively (**Table S1-2**). These parts can be assembled with level 0 parts to construct the expression plasmids of target protein with desired switch variant insertions, through a one-pot reaction to clone individually or in a pooled manner. We first built a hinge library containing 26 hinge designs (1b_001-1b_026) in the context of EKAR’s WW domain phospho-binder (1a_001) and ERK-phosphorylatable substrate (1c_001). We focused on synthetic hinges of different lengths in three major classes: long, flexible GS-rich peptides, alpha helical EAAAK-repeat sequences, and proline-rich AP-repeat sequences. Some natural hinges with known secondary structures were also included (**Fig. 2B**).

**Figure 2.**
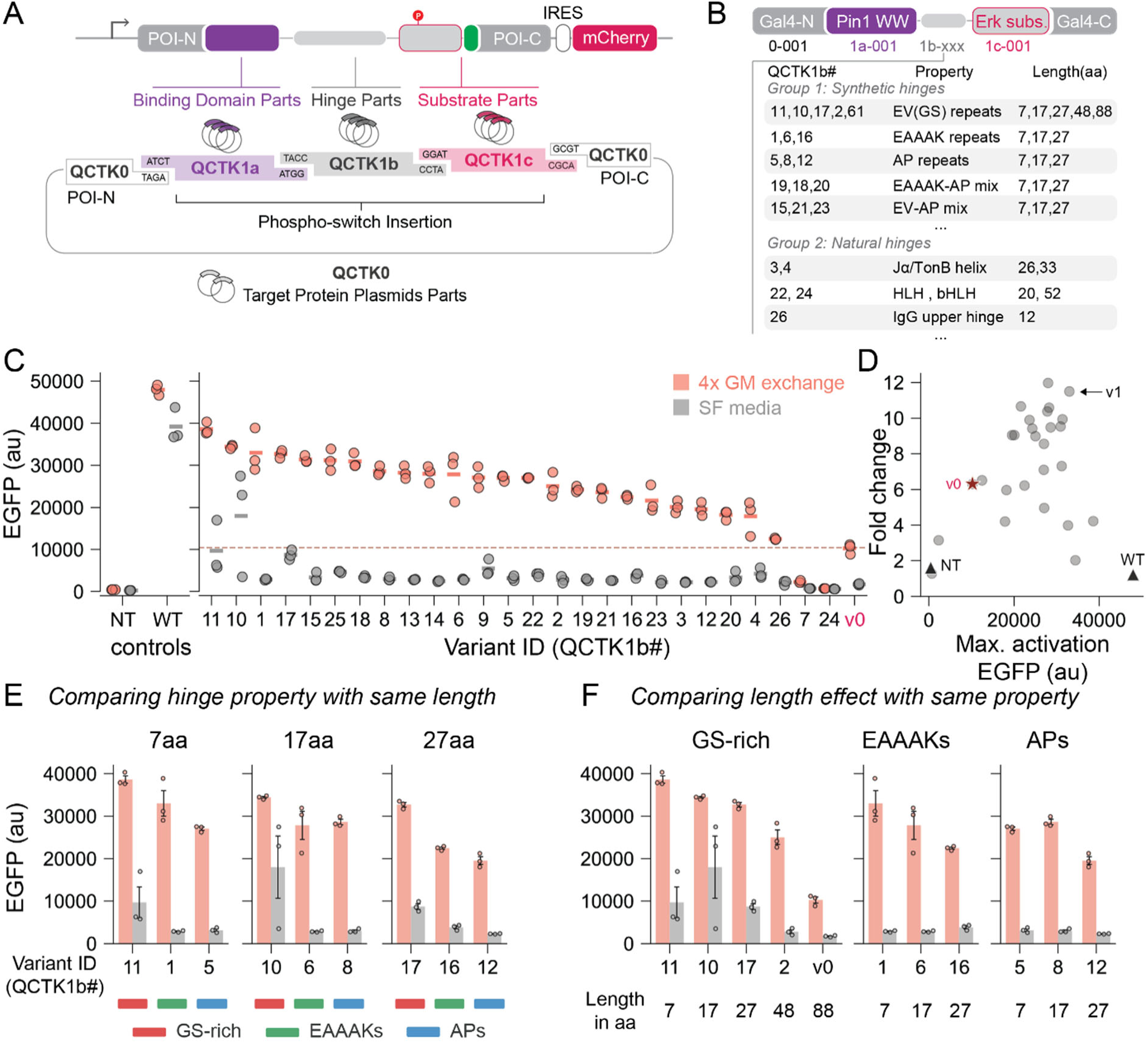
Combinatorial cloning to systematically optimize the hinge in the ERK phospho-switch. (**A**) Schematic of combinatorial Golden-Gate cloning system (QCTK) for rapid phospho-switch variants generation. Three components of the phospho-switch – the phospho-binding domain, hinge, and substrate – are designated as Level 1a, 1b, and 1c parts, respectively. The backbone including the promoter, Gal4 sequences and mCherry marker is designated as Level 0 parts. Assembled plasmids can be immediately transfected and tested in cells. (**B**) Overview of the hinge library. PhosphoGal4 variants were generated for all 26 hinge designs (1b_001-1b_026) with fixed parts for a Pin1 WW domain phospho-binder (1a_001), EKAR biosensor ERK substrate (1c_001) and backbone (0_001). (**C**) Transcriptional responses of all hinge variants in ERK-ON and ERK-OFF conditions. Cells transiently expressing each variant were either kept in serum-free (SF) media or stimulated with four growth media (GM) changes in 30 h before flow cytometry measurements. Bars indicate mean, and 3 replicates are shown as scattered dots. The red dashed line and axis label indicate the performance of the v0 (EKAR) initial design. NT, no transfection; WT, unmodified Gal4-VP64. (**D**) 2D plot of the fold-change and maximum activation for all 26 hinge variants. Red star: v0 (EKAR). Black triangles: no-transfection control (left) and WT-Gal4 control (right). Arrow indicates 1b_001 as an optimal hinge with high fold-change and maximum activation. Hinges are color-coded as: GS-rich (red); EAAAK-repeat helices (green); AP-repeat proline-rich (blue); other sequences (gray). (**E**) Comparison of hinge variants of the same length from different classes. 3 synthetic hinge classes are color-coded as in **D**. Error bars represent mean ± s.e.m (n = 3 replicates). (**F**) Comparison of variants of different lengths from the same class. Error bars represent mean ± s.e.m (n = 3 replicates).

All switch variants were assembled into the Gal4 backbone (Part 0_001) and transiently transfected into HEK293T *5xUAS*-dEGFP cells. Transfected cells were either starved or stimulated with 4 growth media exchanges (**Fig. 1G**) and analyzed by flow cytometry to estimate the phosphorylation-dependent change in transcriptional activity. We identified many hinge variants with stronger maximum activation compared to our initial EKAR sequence (**Fig. 2C**; v0). Plotting the fold-change and maximum activation for all 26 hinges revealed individual variants with high scores on both metrics (**Fig. 2D**), and we chose 1b_001 (7aa EAAAKEA hinge) as the starting point for future phospho-switch engineering (**Fig. 2D**; arrow), terming this variant “phospho-switch v1”.

Further examination of the hinge library responses revealed some design principles. First, it enabled comparison of variants from different classes but with the same length. We found that maximum activity in ERK-ON condition followed the pattern: GS-rich flexible > EAAAK helices > AP proline-rich sequences, consistent with the interpretation that increasing rigidity in the hinge interferes with forming the switch-closed state and reduces maximum Gal4 activity (**Fig. 2E**). However, GS-rich flexible hinge variants tended to exhibit strong background activity, resulting in a low fold change between ERK-OFF and ERK-ON conditions (**Fig. 2E**; hinge classes are color coded as in **Fig. 2D**). In contrast, EAAAK/AP hinges showed a high fold change even at a short length, suggesting that higher hinge rigidity is better able to maintain a switch-open state in the absence of phosphorylation (**Fig. 2E**).

Second, the hinge library allowed us to assess performance of variants from the same class but with different lengths. We observed that increasing GS linker length correlated with lower background activity, as has been observed for FRET biosensors^18^. However, unlike FRET, we found that longer hinges also reduced the maximum EGFP output, leading to a lower overall fold change (**Fig. 2F**). These design principles are consistent with our top hit being a short but relatively rigid variant: a 7 amino acid helical hinge. We fitted a multi-variable regression model with experimental hinge data to predict corresponding switch performance (see **Methods**; **Fig. S3A-C**). In predicting the response of additional computer-generated sequences, the model recapitulated the observed patterns for hinge rigidity and length, suggesting that rigid EAAAK- and AP-repeat sequences exhibited an optimal balance of maximum activation and sensitivity to phosphorylation (**Fig. S3D-F**).

We used AlphaFold3^26^ to predict the structures of both phosphorylated and unphosphorylated ERK phospho-switch v1 in Gal4 (**Fig. S4**), which captured the binding event between the WW domain and phosphorylated ERK substrate, matching previous structural studies of the WW-peptide interaction^27^. We also measured ERK-phosphoGal4v1 performance after inserting additional linker residues between the phospho-switch and Gal4 DNA binding domain, which should weaken allosteric coupling between these domains (**Fig. S5A**). Indeed, we observed decreasing ERK-switchable behavior with increasing linker length, consistent with allosteric coupling that is weakened by longer, more flexible linkers (**Fig. S5B**).

### Improving the substrate site and phospho-binder to generate an optimal phospho-switch

Although our initial phospho-switches exhibited good switching between serum-free and growth media conditions, we found that a non-phosphorylatable mutant switch still retained modest residual EGFP expression in growth media (**Fig. 1H**). We set out to understand and reduce this effect, initially hypothesizing that the residual activity might be due to phosphorylation at additional sites in the substrate peptide that leads to WW domain binding and Gal4 activation.

We constructed a series of phospho-switches harboring mutant substrate peptides (variants of part 1c in the Golden Gate toolkit; **Fig. 3A**) that were designed to eliminate potential ERK-phosphorylatable Ser/Thr residues (Substrates 2-4). We also included a peptide that eliminates all phosphorylatable residues, phospho-mimetic residues, and proline residues, any of which could contribute to WW domain binding (Substrate 5). We found that the presence of the phosphorylatable PRTP motif ensured strong serum-dependent EGFP expression regardless of these additional sequence modifications (**Fig. 3B**), and only Substrate 5 completely abolished residual EGFP expression in growth media (**Fig. 3B**). Based on these data, we designed a final pair of peptides in which the phosphorylatable PRTP or non-phosphorylatable PRAP motif was re-introduced into the otherwise-noninteracting Substrate 5 peptide (**Fig. 3C**). A single Pro-directed phosphorylation site was able to rescue high transcriptional output in growth media, and the corresponding non-phosphorylatable PRAP variant exhibited a 42% reduction in background activity compared to the original ERK-phosphoGal4v1 design (**Fig. 3C**). This new variant, termed “phospho-switch v2”, exhibits a similar maximum activation level, but because of the decrease in background, shows a 1.7-fold increase in dynamic range between PRTP and PRAP variants in growth media compared to the v1 design (**Fig. 3G**).

**Figure 3.**
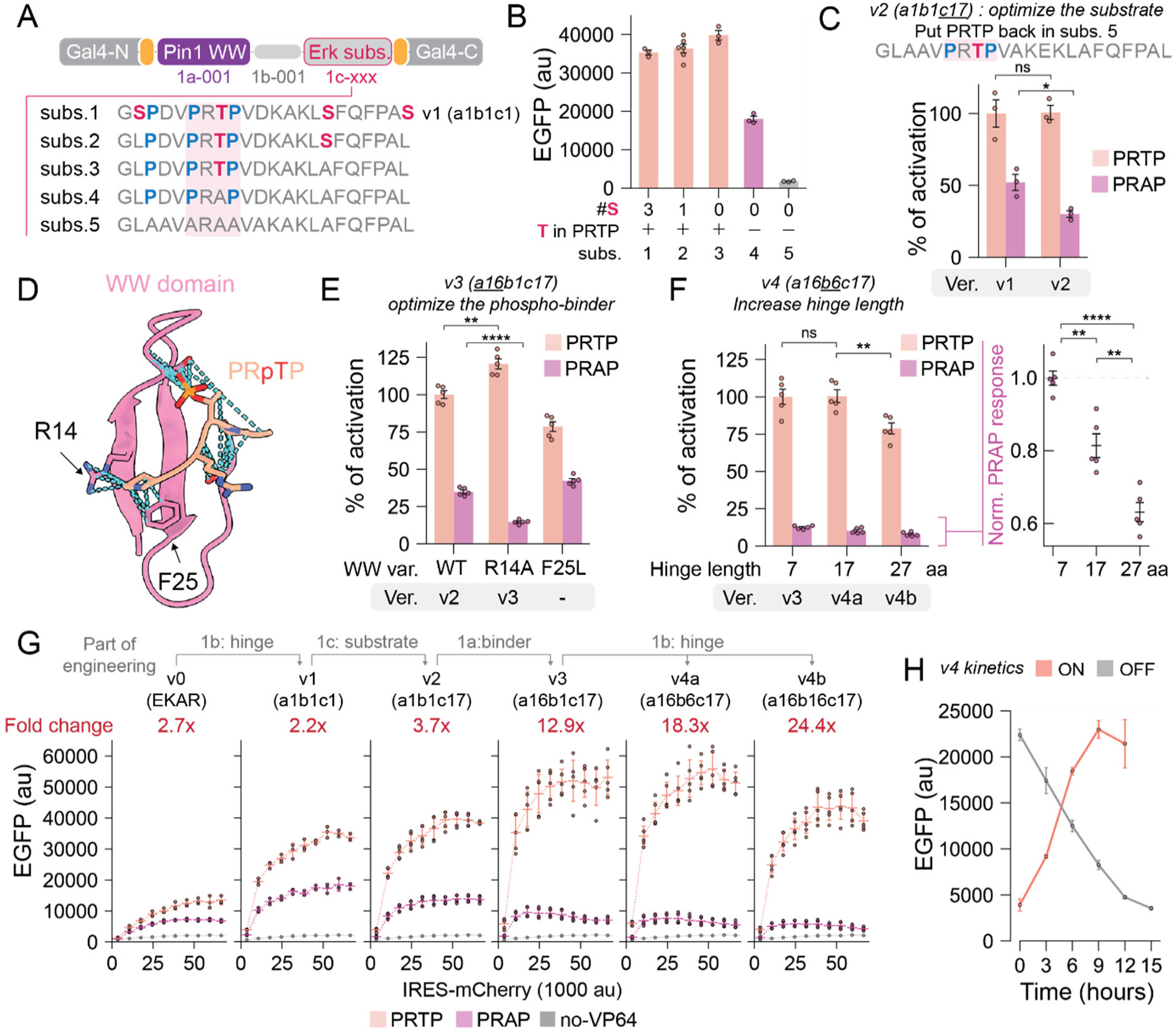
Substrate and phospho-binder optimization for an optimal ERK phospho-switch. (**A**) Overview of mutant substrate peptides. ERK-phosphoGal4v1 (substrate 1) was modified to include substrate peptide variants (substrates 2-5) by mutating phosphorylatable, proline, and negatively charged residues that might contribute to WW domain binding. (**B**) GFP expression of phosphoGal4 variants. Cells transiently expressing variants were stimulated with four growth media changes in 30 h before flow cytometry measurements. Error bars represent mean ± s.e.m (n = 3-6 replicates). (**C**) Reintroducing the ERK-consensus phosphorylation motif (PRTP or non-phosphorylatable PRAP as control) into the noninteracting peptide (subs. 5) increases the switch’s dynamic range. The best-performing variant (a1b1c17) is termed the “v2” phospho-switch. Error bars represent mean ± s.e.m (n = 3 replicates). (**D**) AlphaFold3-predicted model of the Pin1 WW domain (pink) interacting with PRpTP (phosphorylated PRTP peptide, orange). Arrows indicate positions of Arg14 and Phe25. Blue dash lines indicate predicted contacts. (**E**) Characterization of ERK-phosphoGal4v2 variants with WW domain mutations in phosphorylatable (PRTP) and non-phosphorylatable (PRAP) contexts. The resulted optimal variant (a16b1c17) terms as “v3”. Error bars represent mean ± s.e.m (n = 5 replicates). (**F**) Characterization of ERK-phosphoGal4v3 variants with increasing length of helical hinges in phosphorylatable (PRTP) and non-phosphorylatable (PRAP) contexts. The 2 result variants (a16b6c17 and a16b16c17) are termed “v4a” and “v4b”. Error bars represent mean ± s.e.m (n = 5 replicates). (**G**) Overall course of optimization from v0 (EKAR) to v4a/b (a16b6/16c17). Performance of each version is presented as GFP expression curves across a range of TF expression levels. Error bars on the curve represent the mean ± 95% confidence interval (n = 3-5 replicates). Phosphorylatable (PRTP), non-phosphorylatable (PRAP), and a no-VP64 phosphoGal4v3 variant were tested. (**H**) ON/OFF kinetics of ERK-phosphoGal4v4a in stably expressing cells. 24 h after seeding, cells were incubated in serum-free media for OFF kinetics. 18 h post starvation, cells were switched into growth media culture for ON kinetics. Error bars on the curve represent the mean ± s.e.m (n = 3 replicates). For all panels, statistical significance was computed using the two-tailed Student’s t-test. p-value annotation: ns, p > 0.05; *, 0.01 < p <= 0.05; **, 10^-3^ < p <= 0.01; ***, 10^-4^ < p <= 10^-3^; ****, p <= 10^-4^.

We hypothesized that residual activity could be further reduced by targeted mutations in the WW domain to decrease its affinity for the unphosphorylated substrate. Structural and biochemical studies have determined that specific residues in the WW domain, Arg14 and Phe25, make contacts to the first proline residue in the PRTP site and contribute to binding in a phosphorylation-independent manner (**Fig. 3D**)^28,29^. We constructed phosphorylatable and non-phosphorylatable versions of the ERK-phosphoGal4v2 scaffold in which these two residues were mutated to Ala and Leu, respectively^28,29^. We found that while the F25L mutation had an undesirable effect by reducing maximum activation, the R14A variant still drove strong EGFP expression in growth media conditions and exhibited a further 58% reduction in basal activity from a non-phosphorylatable variant compared to ERK-phosphoGal4v2 (**Fig. 3E**). We termed this variant “phospho-switch v3”. As a final improvement, we revised our hinge design to further weaken substrate-WW interactions by increasing the length of the helical hinge from 7 aa to 17 aa or 27 aa, producing “phospho-switch v4a” or “v4b” with further reduction in background activity (**Fig. 3F**).

Over the course of optimization from our phospho-switch v0 to v4a/b designs, the dynamic range of the engineered TF increased from 2.7-fold to 24.4-fold between phosphorylated and unphosphorylated conditions and the maximum activation increased 3.5- to 4.6-fold (**Fig. 3G**; **Fig. S6**). Our EKAR-phosphoGal4v4a reaches a comparable level of gene expression to unmodified Gal4, and the non-phosphorylatable mutant is comparable to a negative control Gal4 construct that lacks the VP64 transactivation domain (**Fig. S6**). We measured the kinetics of EGFP accumulation and loss in response to media changes in *5xUAS*-dEGFP HEK293T cells that stably express the ERK-phosphoGal4v4a construct, observing activation and inactivation with a half-life of ∼6 h (**Fig. 3H**). Finally, we tested the ERK-phosphoGal4v4a system across multiple cellular contexts, including in mouse embryonic stem cells and fibroblasts, where we observed EGFP expression in response to serum stimulation that was eliminated by treatment with a MEK inhibitor (**Fig. S7**). We thus conclude that our ERK phospho-switch v4 is an optimized phosphorylation-actuated protein switch that can be used to achieve potent and reversible control over gene expression in the context of an allosterically-controlled Gal4-VP64 transcription factor.

### Direct and specific activation of ERK-phosphoGal4 using optogenetics

Media switch experiments are a simple method to alter ERK activity for characterizing candidate ERK-phosphoGal4 designs. However, growth media contains complex mixtures of growth factors and other molecules that can alter the activity of many intracellular pathways. We thus set out to characterize our ERK-phosphoGal4 system in response to stimuli that specifically and uniquely activate the ERK cascade. We reasoned that such an approach might also aid in better understanding the residual EGFP expression observed from our v0-v2 non-phosphorylatable switch variants in growth media conditions (**Fig. 1H**; **Fig. 3G**).

We turned to our previously-developed OptoSOS optogenetic system as a method to directly activate the Ras/ERK pathway^30,31^. We engineered *5xUAS*-dEGFP HEK293T cells that stably expressed a blue-light-responsive variant of this OptoSOS system^31,32^, the ERK-phosphoGal4v2 transcription factor, and the ErkKTR-iRFP biosensor (**Fig. 4A-B**). Continuous blue light stimulation drove sustained nuclear export of the ErkKTR biosensor, indicative of sustained ERK activity, and also produced strong EGFP expression in serum-free media (**Fig. 4C**; **Fig. S8**). The light-induced response could be eliminated by treatment with 10 μM of the MEK inhibitor PD 0325901 (**Fig. 4D**). Light-induced EGFP expression could be tuned by varying the intensity or duty cycle of illumination, with increasing EGFP output as the duration of light pulses increased from 2.5’ every 30’ to continuous illumination at 2 mW/cm^2^ (**Fig. 4E-F**). These data indicate that ERK-phosphoGal4 activity can be tuned to intermediate levels by minute-timescale changes in ERK activity, suggesting that the transcription factor is able to rapidly switch between active and inactive states upon phosphorylation in response to these fast changes in illumination conditions.

**Figure 4.**
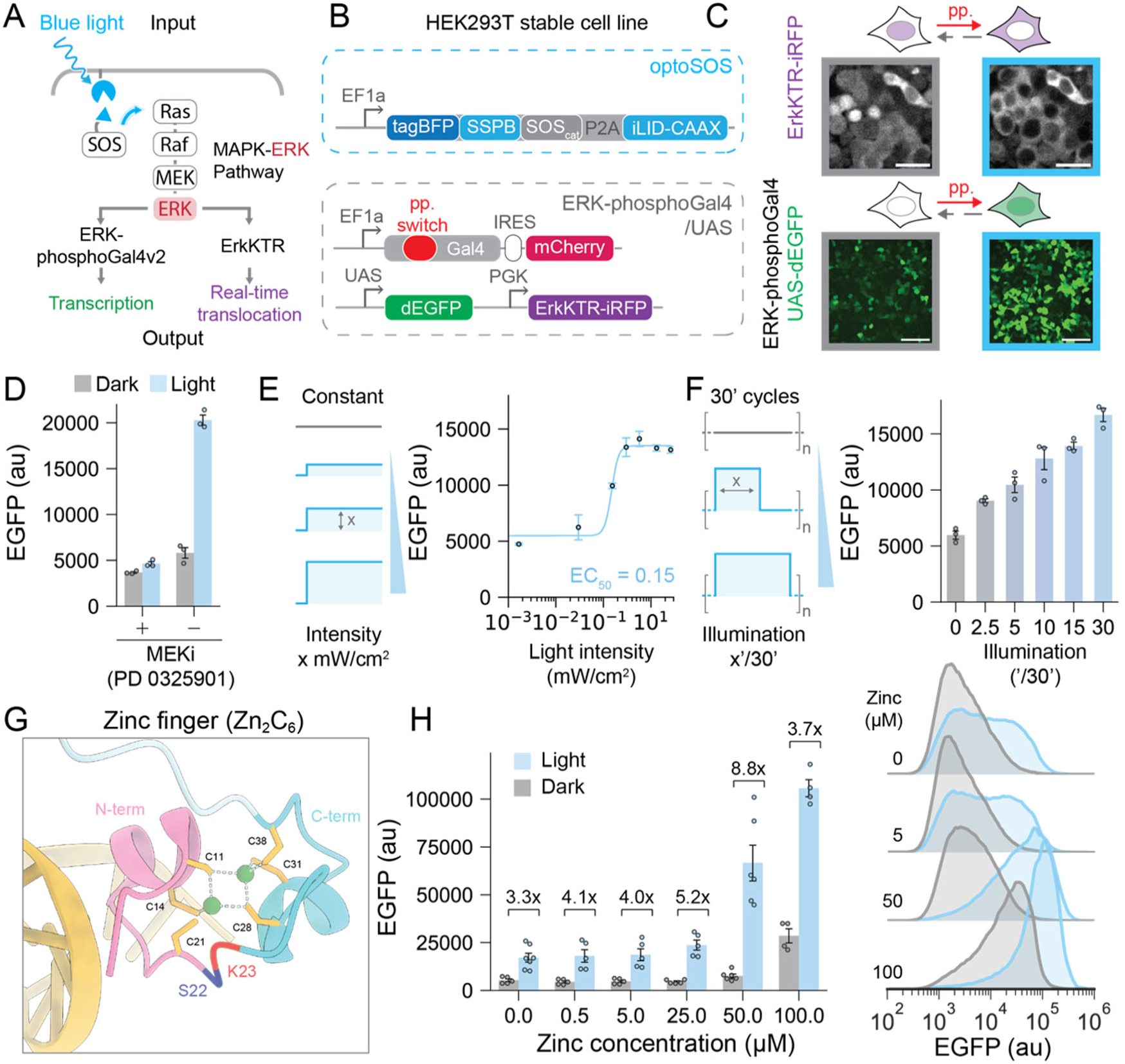
Direct and specific activation of ERK-phosphoGal4 with optogenetics. (**A**) Schematic of optogenetic ERK stimulation and measurements. Blue light induced membrane association of SOS activates the Ras-ERK cascade. ERK activity is directed to the real-time nuclear-to-cytosolic translocation of the ErkKTR biosensor as well as EGFP production through the ERK-phosphoGal4v2/UAS system. (**B**) Overview of the optogenetic HEK293T stable cell line. Cells constitutively express optoSOS, ErkKTR-iRFP, ERK-phosphoGal4v2 and harbor a *5xUAS*-dEGFP reporter. (**C**) Representative images of ErkKTR translocation and ERK-phosphoGal4 induced GFP expression in light and dark. Top panel shows ErkKTR localization before and after 1 h blue light illumination (Scale bar, 25 µm). Bottom panel shows GFP expression before and after 10 h blue light illumination (Scale bar, 100 µm). (**D**) MEK inhibition of ERK-phosphoGal4v2/UAS response to optogenetic ERK stimulation. Cells were incubated in dark or continuous light illumination at 2 mW/cm^2^ for 22 h with or without the presence of 10 μM PD 0325901 and analyzed by flow cytometry. Error bars represent mean ± s.e.m (n = 3 replicates). (**E**) Light-induced GFP expression in response to light intensity levels. Cells were continuously illuminated at various intensities for 22 h and analyzed by flow cytometry. Error bars on the curve represent the mean ± 95% confidence interval (n = 3 replicates). The data is fitted to a Hill curve for estimation of the EC50. (**F**) Light-induced GFP expression in response to illumination duty cycles. Cells were illuminated with various duty cycles ranging from 2.5’ every 30’ to continuous illumination at 2 mW/cm^2^ for 22 h and analyzed by flow cytometry. Error bars represent mean ± s.e.m (n = 3 replicates). (**G**) Crystal structure of Gal4 Zn_2_C_6_ zinc finger (PDB: 3COQ). 6 cysteines (C11, C14, C21, C28, C31, C38) coordinate with 2 zinc ions (green). Conformational switching of a stimulus-sensitive domain between Ser22 (dark blue) and Lys23 (red) is expected to separate the zinc finger into N-terminal (pink) and C-terminal (light blue) halves that may interfere with zinc coordination. (**H**) GFP expression in dark/light conditions in response to media zinc concentration. Cells were incubated in serum-free media supplemented with various levels of ZnCl_2_ in dark or continuous illumination at 2 mW/cm^2^ for 22 h and analyzed by flow cytometry. Error bars represent mean ± s.e.m (n = 4-7 replicates). Histograms show >25000 cells across all replicates.

We also used the simplicity of optogenetic stimulation to further investigate how serum might affect Gal4 activity independently of ERK. The SK22 insertion site is located inside the Zn_2_C_6_ zinc finger (ZF) domain nearby cysteine residues that are essential for Zn^2+^ coordination (**Fig. 4G**)^33^. We hypothesized that the allosteric switch might switch the Gal4 DNA binding domain between zinc-bound and zinc-unbound states. In this model, high zinc ion concentrations could shift the equilibrium to favor the zinc-bound, active Gal4 conformation even in the absence of ERK phosphorylation. Such a model could help explain the difference in activity observed for a non-phosphorylatable variant in serum-free and growth media conditions, as the 1-40 μM levels of Zn^2+^ in tissue culture media is primarily provided by serum^34,35^. To test this model, we used the OptoSOS system to activate ERK in serum-free media supplemented with varying concentrations of ZnCl_2_. Indeed, we observed that high zinc concentrations increased EGFP induction in both illuminated (ERK-ON) and unilluminated (ERK-OFF) conditions (**Fig. 4H**). These data are consistent with a model where high zinc concentrations act in concert with residual WW-peptide binding to shift the equilibrium of ERK-phosphoGal4v2 to the transcriptionally-active conformation. We note that our improved v4 phospho-switches already eliminate serum-induced activity from the non-phosphorylatable PRAP variant, likely by reducing the WW-peptide binding affinity to shift the equilibrium to the switch-open, zinc-unbound state even under growth media conditions.

### ERK-phosphoGal4 outperforms the FOS promoter for sensing ERK activity

Transcriptional biosensors are widely used for assessing signaling pathway activity in high-throughput screens and for reporting on spatiotemporal signaling in multicellular contexts^36–38^. Most transcriptional biosensors are constructed from signaling-responsive endogenous enhancers and promoters^39–41^. These classic reporters suffer from two drawbacks. First, they can lack specificity: endogenous enhancers contain binding sites for many transcription factors and thus their expression is not exclusively determined by the activity of a single kinase. A second challenge is availability, as many endogenous kinases do not drive specific transcriptional responses and thus lack well-defined transcriptional biosensors. We reasoned that one immediate use of our engineered phosphorylation-switchable TF could be as a direct transcriptional biosensor of kinase activity that is more specific for ERK signaling than traditional systems based on endogenous promoters.

We set out to compare our optimized ERK-responsive transcription factor (ERK-phosphoGal4v4a) to a 2.5 kb *c-fos* upstream regulatory sequence (*P_FOS_*). While *P_FOS_* has been widely used to report on ERK activity^42–44^, it contains multiple TF binding sites including cAMP responsive elements (CRE), serum responsive elements (SRE) and AP-1 binding sites, which can serve as targets for a wide range of additional pathways (e.g., PKA, calcium, PKC, and other MAPKs) (**Fig. 5A**)^40,45^. Notably, *P_FOS_* is also widely used as a calcium-responsive promoter of neuronal activity^46,47^. We constructed stable HEK293T cell lines containing either *5xUAS*-dEGFP or *P_FOS_*-dEGFP that each includes a constitutively expressed iRFP marker using PiggyBac-mediated genome integration^48^. Both lines were sorted with similar expressions of iRFP to ensure similar strengths of each reporter. We also stably introduced ERK-phosphoGal4v4 into the *5xUAS*- dEGFP cell line (**Fig. 5A**).

**Figure 5.**
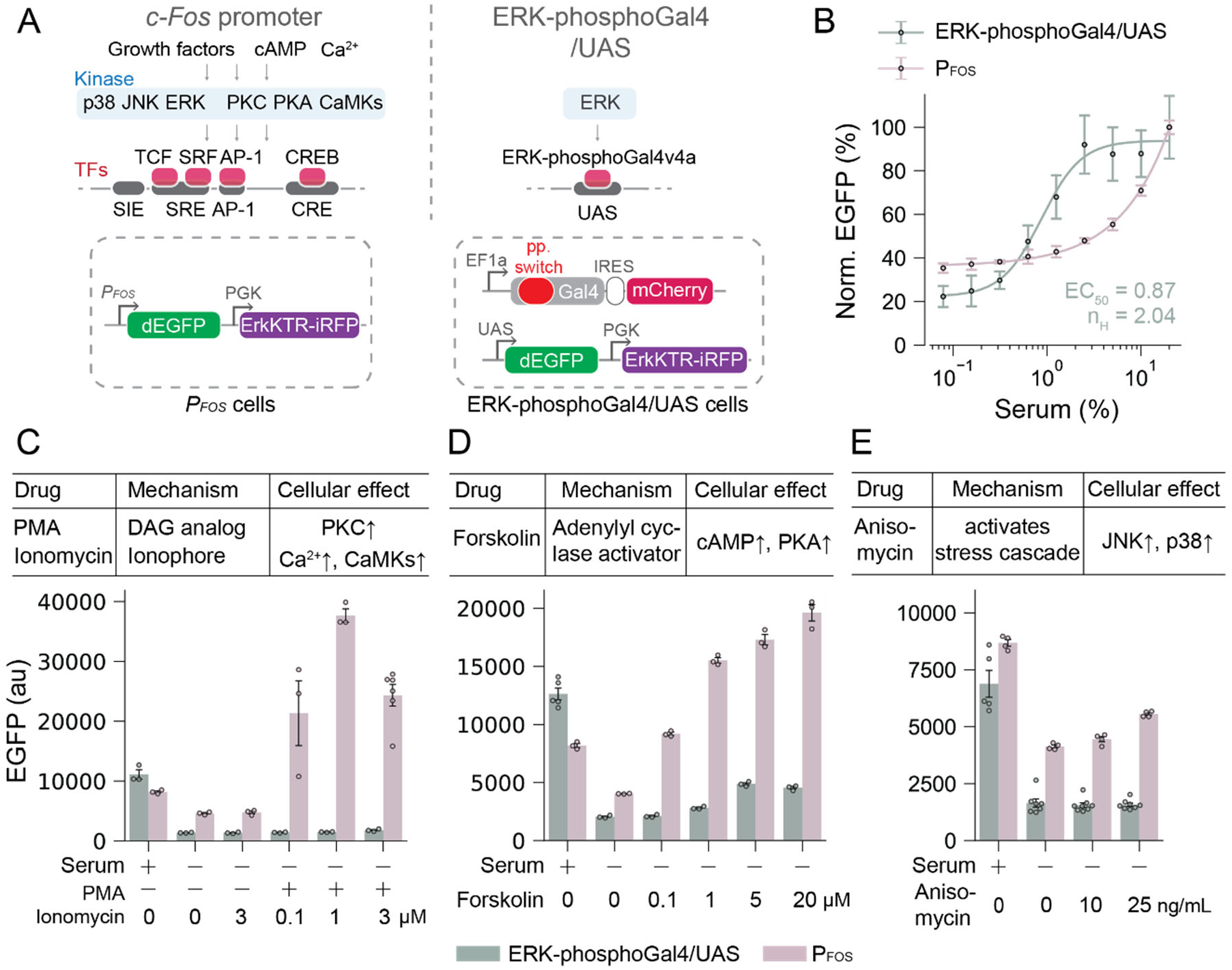
Comparison of ERK-phosphoGal4/UAS and *P_FOS_* promoter. (**A**) Schematic of ERK-to-transcription signal transmission for the *P_FOS_* promoter and the ERK-phosphoGal4v4a/UAS system. Two HEK293T cell lines were constructed to compare both reporting systems at similar strength, using ErkKTR-iRFP as an integration marker. (**B**) Serum dose-response curve for the *P_FOS_* promoter and the ERK-phosphoGal4v4a/UAS system. Cells were incubated in serum-free (SF) media overnight and treated with 0.1-20% serum for 4 h. Each curve is normalized to the maximum GFP expression. Error bars on the curve represent the mean ± 95% confidence interval (n = 3-5 replicates). The ERK-phosphoGal4v4a data is fitted to a Hill curve for estimation of the EC50 and Hill coefficient. (**C-E**): Comparison of *P_FOS_* and ERK-phosphoGal4v4a/UAS systems in cells stimulated with diverse signaling inputs including the PKC/calcium activators PMA and ionomycin (in **C**), the PKA activator forskolin (in **D**), and the JNK/p38 activator anisomycin (in **E**). Cells were pre-starved overnight and incubated in drug conditions for 4 h before flow cytometry measurements. Error bars represent mean ± s.e.m (n = 3-8 replicates). Serum conditions indicate standard growth media (10% FBS). In **C**, 30 nM PMA was used; all other drug conditions are as indicated on the figure.

To assess each cell line’s response to ERK-activating stimuli, we treated them with various concentrations of serum after overnight incubation in serum-free media and analyzed EGFP expression by flow cytometry 4 h post-treatment. These experiments revealed striking differences between ERK-phosphoGal4v4a and the *P_FOS_* promoter (**Fig. 5B**). ERK-phosphoGal4v4 was activated with an EC50 of ∼1% serum, consistent with measurements of serum-induced ERK activity^24^. Gene expression was induced by serum with a Hill coefficient of 2.0 that suggests modest cooperativity. In contrast, the *P_FOS_* response does not plateau even at relatively high serum concentrations of 20%. We also found *P_FOS_* to have high background activity in HEK293 cells when starved, resulting in a low dynamic range compared to ERK-phosphoGal4v4a (**Fig. 5B**). Finally, we compared both systems in the presence of other signaling activators: PMA/ionomycin to activate PKC and calcium signaling, forskolin to activate PKA, and anisomycin to activate the p38 and JNK MAPKs (**Fig. 5C-E**; **Fig. S9**). *P_FOS_* was even more strongly activated by PMA/ionomycin and forskolin than by serum stimulation and was also partially activated by anisomycin treatment. In contrast, ERK-phosphoGal4v4a was insensitive to PMA/ionomycin and anisomycin, with a mild response to forskolin treatment much lower than that observed in response to serum stimulation. Taken together, these results suggest that our engineered ERK-responsive transcription factor, ERK-phosphoGal4v4a, exhibits a well-defined dose response to ERK-activating stimuli and low crosstalk to other intracellular signals, with superior performance relative to *P_FOS_*-based reporters.

### Phospho-switch control is generalizable to other input kinases and effector proteins

Kinase-controlled allosteric switches have the potential to be powerful tools for synthetic biology because of their generalizability to multiple inputs and outputs. At the input level, FRET biosensors have been developed for many kinases, and these phospho-sites could also be incorporated into our optimal switch backbone to rewire input specificity. At the output level, our optimized phospho-switch could also be inserted into other allosterically controlled target proteins. We thus set out to test whether our design could be extended to other input kinases and effector proteins.

We first tested whether the phosphoGal4 design could be rewired to another input kinase (**Fig. 6A**). We turned to the JNK MAPK, a kinase which also has a well-characterized FRET biosensor (JNKAR) but for which transcriptional biosensor options are limited. We assembled a JNK-phosphoGal4 system using the “v2” (wild-type WW phospho-binder) and “v3” (WW^R14A^) architecture, but where the ERK-specific substrate was exchanged for the JNK-specific JNKAR substrate sequence. Each phospho-switch design was inserted into Gal4 at the same SK22 allosteric site (**Fig. 6B**) and transfected into *5xUAS*-dEGFP cells. Indeed, we found that cells expressing a phosphorylatable JNK-phosphoGal4 construct induced EGFP expression in response to the JNK activator anisomycin, with the “v3” design outperforming “v2” (**Fig. 6C**) as we had previously observed for ERK (**Fig. 3G**). EGFP expression varied with anisomycin dose (**Fig. 6D**) over a concentration range previously seen to activate JNK^25^. Importantly, our ERK-phosphoGal4v4a system was not sensitive to anisomycin (**Fig. 5E**), so the observed response is specific to the inclusion of a JNK-targeted substrate. In sum, our phospho-switch design is suitable to be used for multiple input kinases.

**Figure 6.**
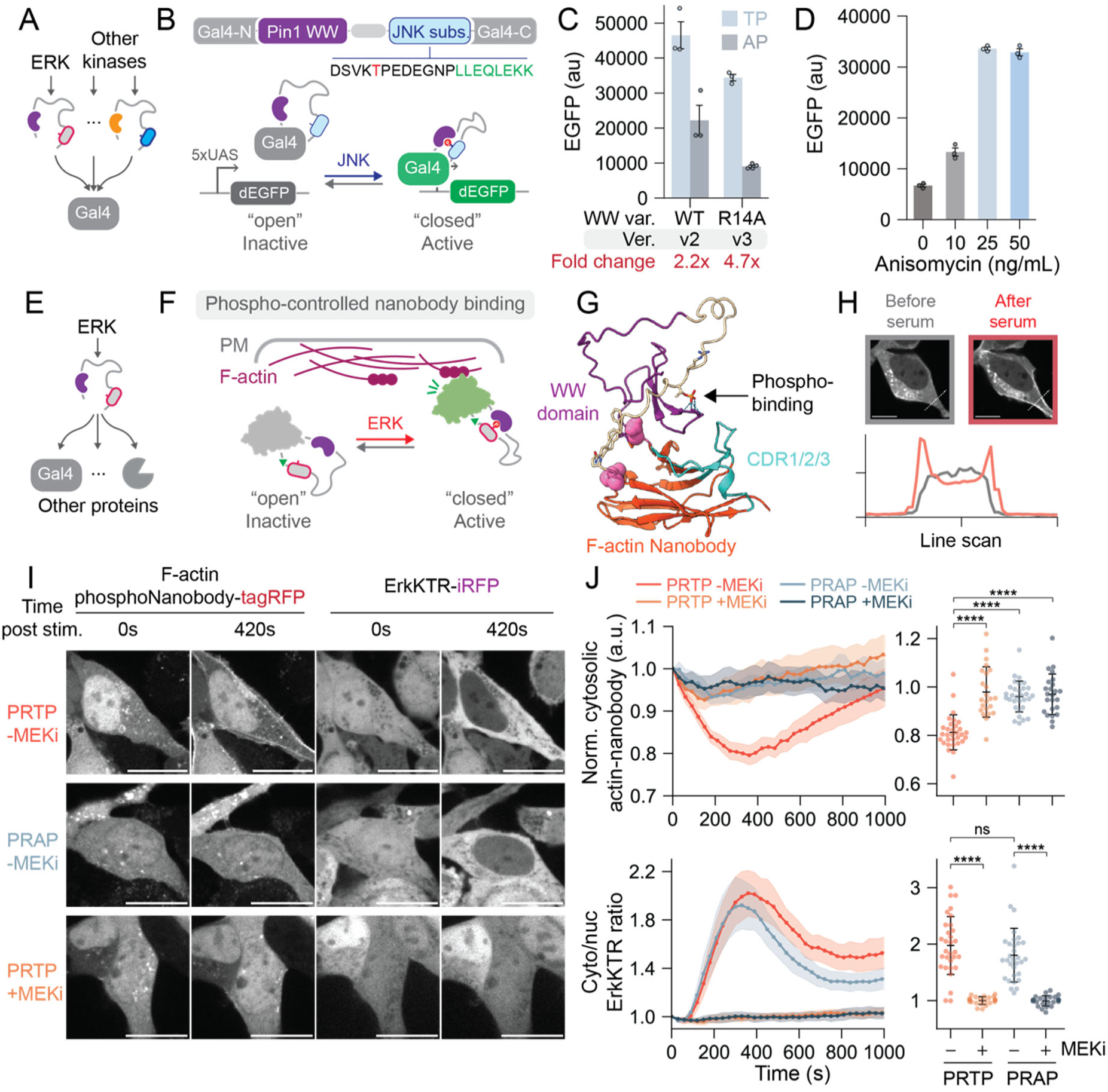
Phospho-switch control is generalizable to other input kinases and effector proteins. (**A**) Generalizing the phospho-switch to accept inputs from additional kinases. (**B**) A JNK-dependent phospho-switch was developed by including a JNK-specific substrate (phosphorylation site in red and docking motif in green). (**C**) Anisomycin-induced GFP expression of JNK-phosphoGal4v2 (harboring the wild-type Pin1 WW domain) and JNK-phosphoGal4v3 (harboring the WW^R14A^ domain) as compared to non-phosphorylatable T-to-A mutants. Cells transiently expressing each JNK-phosphoGal4 variant were incubated in serum-free (SF) media containing 25 ng/mL anisomycin for 24 h. (**D**) Anisomycin dose-response of JNK-phsphoGal4v3. Cells transiently expressing JNK-phosphoGal4v3 were incubated in SF media with various anisomycin concentrations for 24 h. For **C-D**, Error bars represent mean ± s.e.m (n = 3 replicates). (**E**) Generalizing the phospho-switch to control additional output proteins. (**F**) The ERK phospho-switch v3 was inserted into an anti-F-actin nanobody to generate an ERK-phosphoNanobody. (**G**) AlphaFold3-predicted model of the ERK-phosphoNanobody. The ERK phospho-switch was inserted between Asp63 and Gly66 (pink) of the F-actin nanobody (orange; CDR1/2/3 in cyan). Arrow indicates predicted contacts between the phosphorylated threonine and the WW domain (purple). (**H**) Serum-induced ERK-phosphoNanobody binding. Top panels: images of a representative cell before and after 7 min treatment with serum. Dashed lines indicate the position of a line scan for quantifying ERK-phosphoNanobody intensity. Scale bar, 10 μm. Bottom panels: quantification of ERK-phosphoNanobody fluorescence along the line scans before (gray) or after treatment (red) reveals translocation to the cell membrane upon serum stimulation as expected for cortical actin binding. (**I**) Representative ERK-phosphoNanobody and ErkKTR images before and after serum treatment in the presence or absence of 10 μM U0126, a MEK inhibitor. A non-phosphorylatable T-to-A variant was tested as a control. Scale bar, 20 μm. (**J**) Quantification of ERK-phosphoNanobody and ErkKTR response to serum stimulation. Left panels show time traces of cytosolic ERK-phosphoNanobody fluorescence intensity and cytosolic-to-nuclear ErkKTR ratio over time after serum stimulation at 0 sec. Mean trajectories (solid lines) are shown with shading representing the 95% confidence interval. Right panels show single cell responses 7 min after serum treatment. Error bars represent mean ± s.d (For both panels: PRTP -MEKi, n = 31 cells; PRTP +MEKi, n = 23 cells; PRAP -MEKi, n = 31 cells; PRAP +MEKi, n = 24 cells. Cells are quantified from 2-3 independent experiments). Statistical significance is verified with Mann-Whitney-Wilcoxon tests (two-sided with Bonferroni correction). p-value annotation: ns, p > 0.05; *, 0.01 < p <= 0.05; **, 10^-3^ < p <= 0.01; ***, 10^-4^ < p <= 10^-3^; ****, p <= 10^-4^.

We also tested whether the phospho-switch might also extend to other allosterically controllable target proteins (**Fig. 6E**). We turned to OptoNanobodies, nanobodies into which we previously inserted AsLOV2 to confer light-controlled binding to various target proteins: mCherry, EGFP, and endogenous F-actin^49^. We replaced AsLOV2 in the F-actin OptoNanobody with our “v3” variant ERK phospho-switch to generate a candidate ERK-phosphoNanobody (**Fig. 6F**). We inserted the phospho-switch between amino acid positions 63-66, as previously done for AsLOV2 insertion (**Fig. 6G**), fused the construct to TagRFP to monitor its localization, and introduced it into HEK293T cells that also expressed the ErkKTR biosensor to monitor ERK activity^25^. We then assessed localization of the nanobody in growth media stimulated cells for both a phosphorylatable (PRTP) and non-phosphorylatable (PRAP) switch, in the presence or absence of the MEK inhibitor U0126.

Serum stimulation of ERK-phosphoNanobody cells resulted in rapid TagRFP relocalization from the cytosol to the cell periphery (**Fig.6H-I; Mov. S1**), consistent with nanobody binding to cortical F-actin^49^. This effect was not observed in cells expressing a non-phosphorylatable nanobody variant and was also abrogated by treatment with 10 μM of the MEK inhibitor U0126, which blocked ERK activity as measured by redistribution of the ErkKTR biosensor (**Fig. 6I; Mov. S2-4**). We quantified the response by measuring the nanobody’s cytosolic fluorescence intensity, which decreased upon redistribution of the nanobody to cortical F-actin after serum stimulation. Nanobody cytosolic clearance peaked at 6 min after serum stimulation before partially adapting back to baseline within 10-15 min (**Fig. 6J**, upper panel), closely matching the dynamics of ErkKTR translocation (**Fig. 6J**, lower panel). These data suggest that like the ErkKTR biosensor, the ERK-phosphoNanobody rapidly tracks changes in ERK kinase activity. We further tested whether differences in performance could be observed for ERK-phosphoNanobodies harboring “v2” or “v3” versions of the phospho-switch. Indeed, we observed more residual activity in the non-phosphorylatable “v2” phosphoNanobody compared to the “v3” variant with weakened WW binding affinity (**Fig. S10**), suggesting again that the optimized phospho-switch performs well across protein contexts. In sum, these data reveal that our kinase-controlled switch can indeed be generalized across multiple input kinases by swapping phospho-peptides from other FRET biosensors, and can be wired in to control multiple output proteins in which sites of stimulus-controlled allostery have been identified.

## Discussion

Building customizable input/output interfaces with the cell’s endogenous kinase networks is a grand challenge in synthetic biology, with diverse applications in cell engineering, biosensing, and the detection and reversion of disease-associated cellular states. However, our ability to engineer novel phosphorylation-regulated processes in the cell is still limited. Given a kinase and target protein of interest, how might the synthetic biologist draw a new link to trigger an output protein’s activity in response to phosphorylation by an input kinase?

In the current study, we explore this broad question using a combination of recently developed tools. On the output side, engineered allosteric control has been successfully used to design stimulus-responsive variants of enzymes, kinases, transcription factors, and protein binders^15,16,21,49–52^. On the input side, FRET biosensors are available for an increasing number of kinases to convert a phosphorylation event to a conformational change in the biosensor^20^. We set out to link these two emerging approaches by engineering protein switches that convert phosphorylation by a specific kinase into an allosteric change in a target protein’s activity. We find that these kinase-controlled allosteric switches serve as potent, reversible control points for protein function. Our optimized switch – the ERK phospho-switch v4 – can confer up to a 20-fold change in gene expression when inserted at a previously identified allosteric control point in the Gal4 DNA binding domain. The switch design also generalizes to multiple input kinases (ERK and JNK) and output proteins (Gal4 and an F-actin nanobody).

Our study also reveals key design features for the allosteric switch that differ from the original FRET biosensor. For example, we found that the flexible linkers that perform well for FRET make exceptionally poor allosteric switches regardless of length, and that short, rigid linkers combine high maximum activation with low leakiness. We also find that residual binding affinity between the phospho-binding domain and unphosphorylated substrate can be deleterious and identify mutations in both the phospho-binder and substrate that improve the phospho-switch’s dynamic range. Thus, while an ERK-responsive FRET biosensor serves as a useful starting point for switch engineering, optimal phospho-switch performance required substantial additional engineering. It is possible that these advances (e.g. WW domain mutations; short rigid hinge sequences) could be ported back to FRET biosensors to further optimize their performance. Alternatively, some of these design constraints may be unique to the phospho-switch, where a target protein is fused to the switch’s N and C termini and likely exerts forces on them that are not present in the FRET biosensor framework.

Throughout this study we focus on one model system for allosteric control: an engineered zinc finger transcription factor based on the yeast Gal4 DNA binding domain and viral VP64 transactivation domain. The resulting ERK-phosphoGal4v4a transcription factor is able to directly convert ERK kinase activity into up to a 20-fold change in gene expression (**Fig. 3G**). We also show that the sensitivity and specificity of the ERK-phosphoGal4v4a system compares favorably to a classic transcriptional readout, the *FOS* promoter, that is sensitive to many additional signaling cues.

The ability to confer kinase-based regulation over target proteins is likely to be widely useful. We have already developed an ERK-controlled Gal4 transcription factor that could be used to measure kinase activity in complex three-dimensional tissues, where resolution can limit the applicability of existing FRET and translocation-based biosensors^53,54^, or in cases where high-throughput, non-destructive assays are essential, such as FACS-based isolation of rare cell populations with altered kinase signaling^55^. A kinase-directed transcription factor could also be useful as a synthetic biology tool to trigger novel responses to a cell’s signaling state (e.g., by rewiring signaling-hyperactive tumor cells to cytostatic or apoptotic outcomes)^6^, or as a method to add feedback loops to intracellular signaling pathways to alter their signal processing^56^.

This work is only a first step toward a fully generalizable approach in which any kinase could be “wired in” to control any target protein. We have developed an optimized phospho-switch based on the WW phosphorylation-binding domain, but additional binding domains (e.g. FHA1) may be required to broaden the range of kinases and substrates that can be used^57^. Phospho-tyrosine signaling could also be used within a phospho-switch in a similar manner, using the specific interaction between certain phosphotyrosine peptides and SH2 domains^14,58^. Finally, it will be essential to broaden the set of target proteins with potent sites for allosteric control so that a diverse set of cellular outputs will be accessible to novel kinase-based control^15,21,49,51^. These tools and others may help to usher in a new era of kinase-controlled transcriptional responses and synthetic post-translational intracellular logic circuits.

## Methods

### Plasmid construction

Important plasmids used in this study are listed in Supplementary Table 3. pQC060 was constructed by cloning the wildtype Gal4-VP64 expression cassette from pHR_SFFV_Gal4_VP64_IRES_mCherry (Addgene, #203909) into a PiggyBac backbone. EKAR (88aa hinge version) was amplified from a home-made modified version of pPBJ-EKAR-EV-NES (a gift from University of California-Davis, Albeck lab) with the 116aa hinge shortened to 88aa. pQC061 was constructed by inserting the EKAR amplicon into pQC060 at the Ser22/Lys23 allosteric site. pQC171/235 was constructed by inserting and replacing the original AsLOV2 insertion with ERK phospho-switch v2/v3 in pHR_SFFV_Actin_optoNB_tagRFP (Addgene, #159595). Phosphorylation site T-to-A mutations were introduced to pQC061 (this study, EKAR-Gal4) via whole backbone PCR using primers containing the target mutation to generate pQC086. Mutations were introduced using the same procedure on 0_001_v2_a1b1c4 (this study, JNK-phosphoGal4v2), pQC256 (this study, JNK-phosphoGal4v3) and pQC171/235 (this study, F-actin ERK-phosphoNanobodies) to generate pQC188, pQC257 and pQC172/pQC236. Other plasmids for cell line generation (this study, pQC131, pQC136 and pQC137) were constructed with a PiggyBac backbone. The no-VP64 control plasmid, pQC238 was constructed by whole backbone PCR to exclude the VP64 in 0_001_v2_a16b1c17 (this study). All plasmids discussed in this section were constructed through In-Fusion assembly (TaKaRa, #638911). All other plasmids not mentioned in this section were generated with Golden Gate Toolkit (QCTK) and will be separately discussed in following sections.

### Golden Gate Toolkit (QCTK) Development

Golden Gate Toolkit for fast phospho-switch generation (QCTK) includes Level 0, 1a, 1b and 1c parts (**Fig. 2A**). To generate Level 0 parts, target protein expression vectors were first cloned with desired backbones, promoter sequences and expression markers. For example, a wild type Gal4-VP64 (as target protein) was first cloned into a PiggyBac integration vector with *P_EF1a_* as promoter and IRES-mCherry as expression marker. The cloned expression vectors were opened at the allosteric insertion site with BsaI overhangs by PCR amplification. A bacterial EGFP expression cassette (BBa_J72163_P_GlpT__sfGFP) was amplified from pYTK001 (Addgene, #65108) with corresponding BsaI overhangs. Digestion with BsaI_HFv2 (New England Biolabs, #R3733S) and ligation with T4 DNA Ligase (New England Biolabs, #M0202S) was then conducted to insert the bacterial EGFP marker into the target protein expression vector and leave BsaI sites on both ends of the EGFP marker. All Level 0 parts are carbenicillin-resistant and successful transformants express EGFP on LB agar plates with carbenicillin (Gold Biotechnology, #C-103-5).

To generate Level 1 parts, pYTK001 was amplified as backbones with 1a, 1b or 1c BsmBI overhangs. Desired insert sequences were generated by PCR amplification, oligonucleotide annealing or direct DNA fragment synthesis with corresponding 1a, 1b or 1c BsmBI overhangs. Digestion with BsmBI_v2 (NEB, #R0739S) and ligation with T4 DNA Ligase was conducted to replace the bacterial EGFP marker in pYTK001 with target inserts. All Level 1 parts are chloramphenicol-resistant and successful transformants were picked by EGFP counter-selection on LB agar plates with chloramphenicol (Gold Biotechnology, #C-105-5). Standardized overhang sequences for Level 0, 1a, 1b and 1c parts are shown in **Fig. 2A**. See Supplementary Table 1 QCTK1a/c parts) and 2 (QCTK1b parts) for lists of all QCTK parts used in this study.

### Variant assembly with QCTK

Golden Gate assembly was performed in a one-pot reaction with Level 0, 1a, 1b and 1c parts. Reaction mixes to assemble variant expression plasmids were prepared individually with all 4 parts (50-100 ng for each part), 0.5 μL BsaI_HFv2, 0.5 μL T4 DNA Ligase, 1 μL T4 DNA Ligase Reaction Buffer (New England Biolabs, #M0202S) to a total volume of 6.5 μL. Reaction mix was incubated for 50x digestion/ligation cycles (2 mins at 37 °C followed by 4 mins at 16 °C). Additional digestion for 10 mins at 37 °C followed by heat inactivation for 10 mins at 80 °C was conducted after the digestion/ligation cycles. Reaction mixes were then transformed into Stellar™ Competent Cells (TaKaRa, 636766) and an aliquot of the recovery culture was spread on an LB agar plate with carbenicillin. Successful transformants were picked by counter-selection on EGFP expression (undigested Level 0 part contains the bacterial EGFP expression cassette and undigested Level 1a-c parts are chloramphenicol-resistant). See Supplementary Table 3 for all important plasmids used in this study generated by QCTK Golden Gate assembly.

### Cell culture

Lenti-X HEK293T cells and NIH 3T3 cells were maintained in DMEM (Gibco, #11995073) with 10% FBS (Fetal Bovine Serum; R&D Systems, #S11150), 1% penicillin/streptomycin (Gibco, #15140-122), 2mM L-Glutamine (Gibco, #25030-081) throughout all experiments. For stem cell cultures, basal growth medium was made with GMEM (Millipore Sigma, G6148), 10% ESC-qualified fetal bovine serum (R&D Systems, S10250), 1×GlutaMAX (Gibco, 35050-061), 1×MEM non-essential amino acids (Gibco, 11140-470050), 1 mM sodium pyruvate (Gibco, 11360-070), 100 μM 2-mercaptoethanol (Gibco, 47121985-023) and 1% penicillin/streptomycin. Mouse embryonic stem cell (mESC) line, E14tg2a (ATCC, CRL-1821), was maintained in 0.1% gelatin coated 25 cm^2^ tissue culture flasks and cultured in 2i+LIF media comprising basal growth medium further supplemented with 1,000 units/ml^−1^ LIF (Millipore Sigma, ESG1107), 2 μM PD0325901 (Tocris, 4192) and 3 μM CHIR (Tocris, 4423). Cells were verified by STR profiling (ATCC). Cells were not cultured in proximity to commonly misidentified cell lines.

### Cell transient transfection and ERK stimulation with conditioned media

HEK293T reporter cells were seeded in 48-well plates. 24 hours post seeding, cells were transfected at 60%-80% confluency. Transfection mixtures for each well were prepared in 15 µL Gibco OptiMEM (Gibco, 31985070) containing 2.5 µL FuGENE HD (Promega, #E2311) and 150 ng vector plasmid. 24 hours post transfection, culture media was switched to serum-free (SF) media (DMEM with 1% penicillin/streptomycin and 0.00476 mg/mL HEPES (Sigma-Aldrich, H4034)) or fresh growth media (GM, recipe mentioned in Cell culture section). For reactivation, media was replaced with GM at a defined time after first media switch with serum-free media (in **Fig. 1F, 3H**). For delivering multiple pulses of ERK activation with media change, 75% of the existing media in wells were replaced with fresh GM at 6, 18, 24 h after the first media change (0 h). All ERK-ON experiments in **Fig. 1H, 2 and 3** were done with 4 GM changes in 30 h. After treatment, cells were then detached with 1 min TrypLE (Gibco, 12604–013) treatment at 37 °C and resuspended into a single-cell solution for flow cytometry analysis. For the hinge variant screen, variants were tested in batches with 3 controls in each batch, including a no-transfection control, a WT-Gal4 control and an ERK-phosphoGal4 variant (0_001_a1b1c1) as reference. Batch correction was done individually for ERK-ON and ERK-OFF conditions using the controls and normalized to the first batch.

### Cell line generation

Engineered HEK293T stable cell lines were generated using the piggyBac random integration system. A chassis cell line was first cultured to 60%-80% confluency in 12-well plates. Transfection mixtures were prepared in 45 µl OptiMEM containing 4 µl FuGENE HD, 800 ng vector plasmid and 250 ng PBase ‘helper’ plasmid (System Biosciences). After preparation, the transfection mixtures were equilibrated at room temperature for 15 min and then added dropwise to the chassis cells. Cultures were propagated for at least four days following transfection. The cells were then detached and resuspended into a single-cell solution for fluorescence-activated cell sorting (FACS) using a Sony SH800S sorter. Sorted fluorescence-positive cells are collected and cultured for experiments.

### Flow cytometry analysis and fluorescence-activated cell sorting (FACS)

Flow cytometry analysis or FACS was conducted with a Sony SH800S equipped with Sony 100 μm Sorting Chip. Live, single cells were gated using FSC-A and SSC-A (for live cells) and FSC-A and FSC-H (for single cells). EGFP was excited with a 487.5 nm laser, and emitted light was collected after passing through 525/50 nm band pass filters. mCherry was excited with a 561 nm laser, and emitted light was collected after passing through 600/60 nm band pass filters. iRFP was excited with a 639 nm laser, and emitted light was collected after passing through 720/60 nm band pass filters. Gain configuration and fluorescence compensation was performed following manufacturer instruction. For FACS, at least 100,000 cells were sorted and plated on a new 6-well plate. Detailed strategies of gating, analysis and visualization of flow cytometry data are shown in Fig. S1.

### Fold change calculation

Cells were first gated for IRES-mCherry expression level between 5×10^4^ to 1.25×10^5^ and fold change of the phosphoGal4s were calculated as:

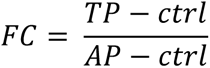

where FC is the fold change reported, TP is the response of phosphoGal4s with phosphorylatable substrate; AP is the response of phosphoGal4s with non-phosphorylatable substrate and ctrl is the response of the control phosphoGal4 lacking VP64.

### Multi-variable linear regression model to predict hinge performance

A multi-variable linear regression model was built to use five selected hinge sequence features (length, A%, GS%, EK%, P%) as input variables and output the hinge performance (include maximum activation in ERK-ON condition and background activation in ERK-OFF condition). The model was trained using experimentally tested hinges. Features extracted from experimentally tested hinges were tested negative for multicollinearity with all variance inflation factors (VIF) < 5. The trained model was then used to predict the performance of a library of > 6000 *in silico* generated hinge sequences. The library includes GS-rich flexible hinges, EAAAK-repeat alpha helical hinges, AP-repeat proline-rich hinges and computationally random mutated hinges at various lengths. Details of the model are shown in **Fig. S3**.

### Optogenetic activation of ERK-phosphoGal4 with optoWELL and flow cytometry

Polyclonal HEK293T cells stably expressing both the optoSOS system and the ERK-phosphoGal4v2/UAS system (**Fig. 4A-B**) were seeded in glass-bottom black 24-well plates (Cellvis, P24-1.5H-N) and kept in dark in the growth media. 24 hours after seeding, culture media was changed to serum-free (SF) media and the plate was then placed on a light-stimulation device (opto biolabs, optoWELL). Each well can be illuminated with 450 nm blue light at a defined intensity and duty cycle profile. Light intensity of the device (in mW/cm^2^) is measured with a power meter (Thorlabs, PM100D). After 22 h of stimulation, cells were detached and analyzed with Sony SH800S Cell Sorter. For MEK inhibition experiments, PD 0325901 (Tocris, 4192) was diluted in serum-free media at a final concentration of 10 μM and cells were kept in PD 0325901-contaning serum-free media throughout 22 h of stimulation. To supplement the system with zinc, additional ZnCl_2_ (Sigma-Aldrich, 208086) was added to the serum-free media.

### Signaling pathway stimulation

Polyclonal HEK293T cell lines harboring *P_FOS_*-dEGFP or ERK-phosphoGal4v4/UAS-dEGFP reporting system were seeded in 48 well plates. 24 h after seeding, cells were incubated in 200 uL serum-free (SF) media overnight (∼18 hrs.). After overnight starvation, drugs used for various stimulation are pre-diluted in serum-free media in 2x concentration. 100 uL previously in-well media was removed and replaced with 2x stimulation media. 4 h after stimulation, cells were detached and analyzed with Sony SH800S Cell Sorter. Following drugs were used in the panels: PMA (Phorbol 12-myristate 13-acetate; Sigma-Aldrich, P8139), ionomycin (Sigma-Aldrich, 407950), forskolin (Calbiochem, 34-428-25MG) and anisomycin (Sigma-Aldrich, A9789).

### Dose-response curve fitting

Dose-response curve in Fig. 4E/5B was fitted with a four-parameter logistic (4PL) model. The model was defined as:

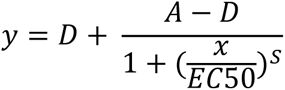

where D is the maximum response, A is the minimum response, EC50 is the half-maximal effective concentration and S is Hill’s slope of the curve. The concentration or intensity of the input and the mean response were used to fit the model.

### Live-cell imaging

Cells were kept at 37 °C with 5% CO_2_ for the duration of all imaging experiments. 50 μL of mineral oil was pipetted onto the wells right before placing onto the scope to prevent media from evaporating. Imaging was done using Nikon Eclipse Ti microscope with a Prior linear motorized stage, a Yokogawa CSU-X1 spinning disk, an Agilent laser line module containing 405, 488, 561 and 650 nm lasers, an iXon DU897 EMCCD camera and objective lenses as specified for individual experiments in the following sections.

### Optogenetic activation of ERK-phosphoGal4 with live-cell imaging

Polyclonal engineered HEK293T cells stably expressing both the optoSOS system and the ERK-phosphoGal4v2/UAS system (**Fig. 4A-B**) were seeded in glass-bottom black 96-well plates (Cellvis, P96-1.5H-N) and kept in dark in the growth media. 24 h after seeding, culture media was changed to serum-free (SF) media and the cells were kept in starvation and in dark for ∼24 h to ensure low baseline EGFP before imaging and stimulation on microscope. Cells were then imaged with previous specified microscope configuration with a 20x air objective lens. A 450 nm LED light source was used for photoexcitation with blue light, which was delivered through a Polygon400 digital micro-mirror device (DMD, Mightex Systems). The LED power was adjusted to 75% to deliver ∼15 mW/cm^2^ blue light at the sample plane, as measured by with a PM100D power meter (Thorlabs). Light was delivered as 1 s pulses of illumination every 1.5 mins throughout 17 h of imaging. ErkKTR-iRFP was captured every 3 mins, EGFP induction and IRES-mCherry expression were captured every 1 h throughout the entire imaging session.

For ErkKTR quantification in this experiment, cells were segmented on whole cell fluorescence in the mCherry channel using Cellpose3^59^. tagBFP, EGFP, mCherry and iRFP signal can be measured inside individual cell masks. Inside the segmented mask for an individual cell, the tagBFP signal was thresholded to generate a cytosolic region mask due to the natural cytosolic localization of the tagBFP-SSPB-SOScat protein. Whole cell masks can then be further divided into cytosolic masks and nucleus masks. ErkKTR-iRFP intensity was measured separately in both regions for each cell.

### Live-cell imaging for the F-actin ERK-phosphoNanobody

Monoclonal HEK293T cells expressing NLS-tagBFP and ErkKTR-iRFP were seeded at low cell density in fibronectin-coated 96 well glass-bottom plates (fibronectin coating concentration, 5 μg/cm^2^). 24 h after seeding, cells were transfected with F-actin ERK-phosphoNanobody expression plasmids using FuGENE HD (same procedure as previously described transfections). 24 h after transfection, cells were starved overnight (>12 hours) with serum-free media and then imaged with previous specified microscope configuration with a 60x oil immersion objective lens. For MEK inhibition conditions, cells were pre-treated with U0126 (catalog) at a final concentration of 10 μM for at least 30 mins prior serum stimulation. To apply the MEK inhibitor, 1 μL 10 mM U0126 stock was pre-diluted in 19 μL serum-free media and 2 μL from the dilution was added in wells with a total media volume of 100 μL at least 30 mins before imaging. Images of iRFP (on ErkKTR) and tagRFP (on F-actin ERK-phosphoNanobody) were taken every 30 s for >20 mins. 2.5 mins after the start of imaging, 2.5 μL of FBS was added directly into wells to activate ERK signaling.

## Author Contributions

Conceptualization, Q.C., J.E.T.; Methodology, Q.C., J.E.T.; Investigation, Q.C.; Funding, J.E.T.; Writing and Editing, Q.C., J.E.T.; Supervision, J.E.T.

## Acknowledgements

We thank all members of the Toettcher lab for helpful discussions and Beatrice Ramm, Emily Kolenbrander Ho, and Long Nguyen for comments on the manuscript. This work was supported by the National Institutes of Health grant R01GM144362 and NSF RECODE grant 2134935 (to J.E.T.).

