## Supplementary Information for "Design and optimization of a kinase-controlled allosteric switch"

**Table of Contents**

Supplementary Figure 1. Example flow cytometry data gating strategy for phosphoGal4/UAS experiments.

Supplementary Figure 2. ERK activity adaptation and reactivation in HEK293T cells.

Supplementary Figure 3. Multi-variable linear regression model to recapitulate the hinge design rules.

Supplementary Figure 4. AlphaFold3 predicted structural models of ERK-phosphoGal4v1.

Supplementary Figure 5 Allosteric un-coupling with ERK-phosphoGal4v1

Supplementary Figure 6. Course of ERK phospho-switch optimization

Supplementary Figure 7. ERK activity biosensing with ERK-phosphoGal4v4a/UAS system in other cell lines

Supplementary Figure 8. Optogenetic activation of ERK over long time periods

Supplementary Figure 9. Histograms for ERK-phosphoGal4v4a/UAS and $P_{FOS}$ promoter comparison

Supplementary Figure 10. Comparison of F-actin ERK-phosphoNanobody with v2 or v3 ERK phospho-switches.

Supplementary Table 1. List of QCTK1b parts

Supplementary Table 2. List of QCTK1a/c parts

Supplementary Table 3. List of important plasmids used in this study

Supplementary Table 4. List of important phospho-switch/phospho-protein sequences in this study

Supplementary Movie 1. F-actin ERK-phosphoNanobodyv3 (PRTP) and ErkKTR responses to serum stimulation with no MEK inhibition

Supplementary Movie 2. F-actin ERK-phosphoNanobodyv3 (PRTP) and ErkKTR responses to serum stimulation with MEK inhibition

Supplementary Movie 3. F-actin ERK-phosphoNanobodyv3 (PRAP) and ErkKTR responses to serum stimulation with no MEK inhibition

Supplementary Movie 4. F-actin ERK-phosphoNanobodyv3 (PRAP) and ErkKTR responses to serum stimulation with MEK inhibition

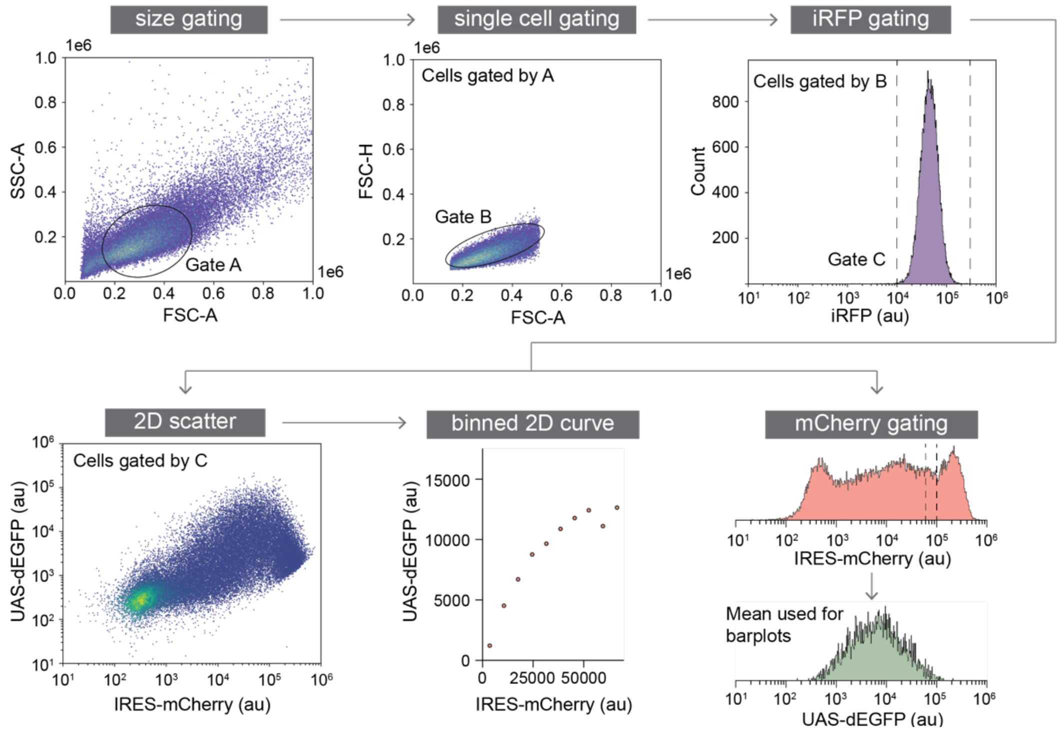


**Supplementary Figure 1. Example flow cytometry data gating strategy for phosphoGal4/UAS experiments.** PhosphoGal4 were transiently expressed and tested in a clonal *5xUAS*-dEGFP HEK293T cell line (**Fig. 1E**). Samples were analyzed for GFP induction with flow cytometry. First, cells are gated for proper single cells with Gate A/B using the FSC and SSC channels (size/single cell gating). Next, Gate C selects cells expressing the integration marker ($P_{PGK}$-iRFP) on the same construct with *5xUAS*-dEGFP (iRFP linear gate:$1\times10^4$-$2\times10^5$). Two analysis pipelines are used for data presentation. (1) Gate C gated cells are then gated for IRES-mCherry expression (mCherry linear gate: $5\times10^4$-$6\times10^4$ or otherwise specified). mCherry-gated cells are used to plot GFP histograms or calculate the mean GFP value for bar graphs. (2) Gate C gated cells are plotted for mCherry v.s. GFP 2D scatter plots. For visualization, we bin cells on IRES-mCherry expression level and generate the IRES-mCherry v.s. GFP curve to show the transcriptional activity cross all expression levels of phosphoGal4.

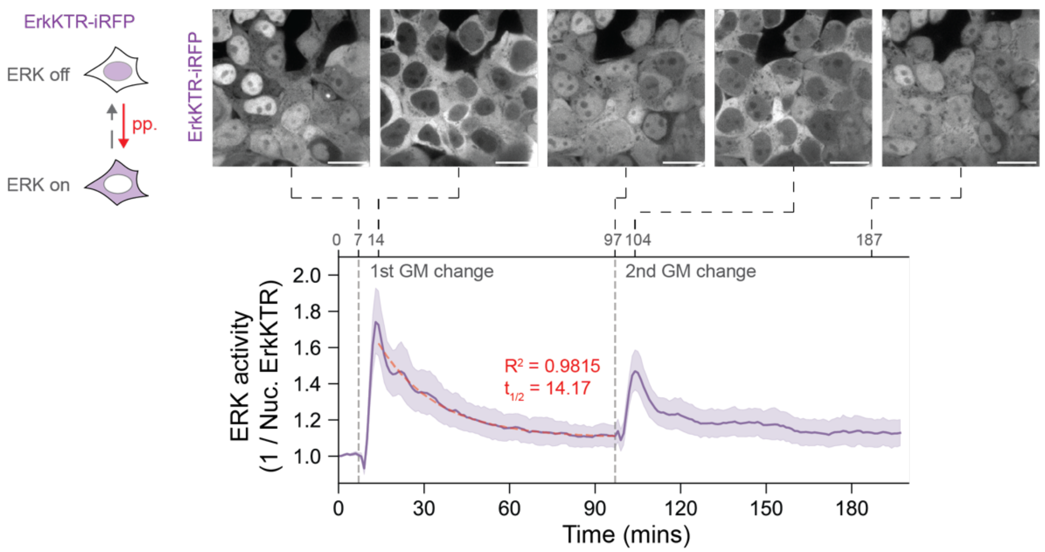


**Supplementary Figure 2. ERK activity adaptation and ERK reactivation in HEK293 cells.** HEK293T cells expressing live-cell ERK biosensor (ErkKTR) were stimulated with growth media (GM, containing 10% FBS). Two GM changes were done following the same procedure of GM change in **Fig. 1G** during imaging. Nucleus ErkKTR clearance was measured to represent ERK activity (n=24). ERK activity peaks 7 mins post GM change and quickly decays. By fitting ERK activity curve to an exponential decay model, the adaptation shows a $t_{1/2}$ of 14.17 mins. ERK activity can also be reactivated by the 2$^{nd}$ GM change and then show similar adaptation. Representative images are shown at the time of 1$^{st}$/2$^{nd}$ GM change, ERK activity peaks and 90 mins post GM change. Scale bars, 20 µm.

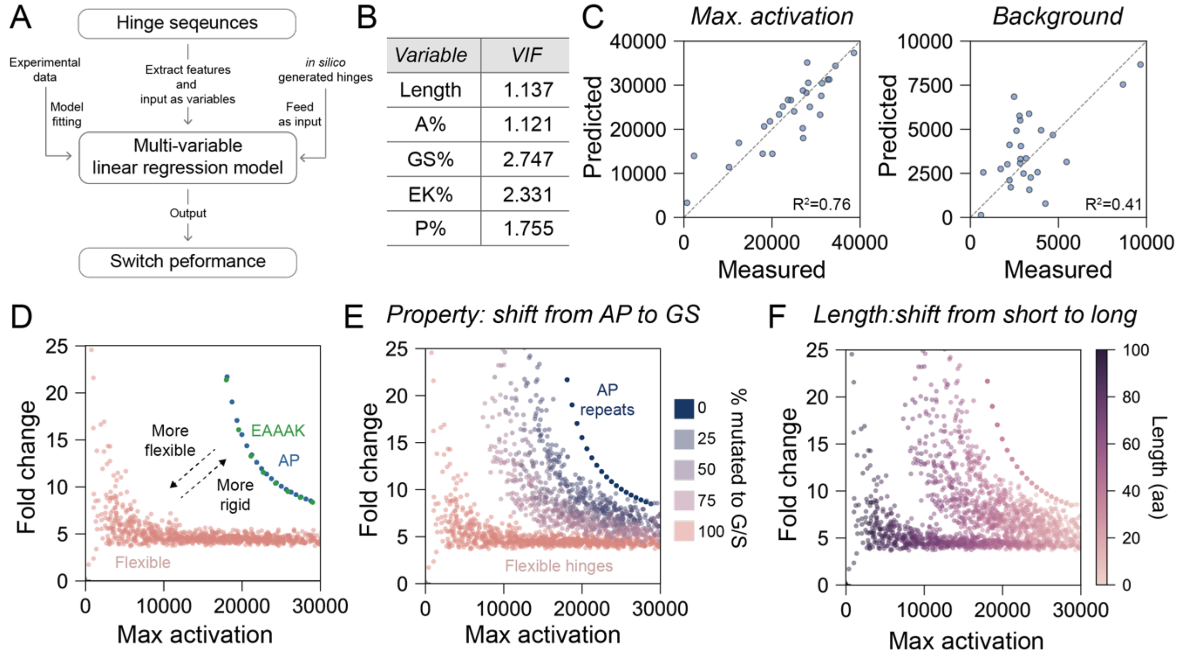


**Supplementary Figure 3. Multi-variable linear regression model to recapitulate the hinge design rules.** (**A**) Features were extracted from hinge sequences and input to a multi-variable linear regression model for predicting phospho-switch variants performance. Sparse experimental data were used for model fitting. Fitted model was then used to predict performance of > 6000 *in silico* generated hinge sequences. (**B**) 5 selected hinge sequence features (length, A%, GS%, EK%, P%) were used as variables and tested for multi-collinearity with variance inflation factor analysis (VIF). All variables show VIF < 5. (**C**) Linear regression model was trained with experimental data to use hinge features to predict maximum activation in ERK-ON conditions and background activation in ERK-OFF conditions of phosphoGal4 variants. (**D**) Computationally generated hinge sequences (GS-rich flexible hinges, EAAAK-repeat alpha helical hinges, and AP-repeat proline-rich hinges) were fed into the model to generate a 2D plot (as in **Fig. 2D**) and the result recapitulates the pattern on hinge rigidity. (**E**) Starting from rigid AP-repeat proline-rich hinges, different percentages of the amino acids were computationally mutated to G/S and increasing percentages of mutations gradually shifts the performance of phosphoGal4 towards those with flexible hinges. (**F**) Model prediction recapitulates the pattern on hinge length. Increasing hinge length correlates with lower maximum activation and higher fold change.

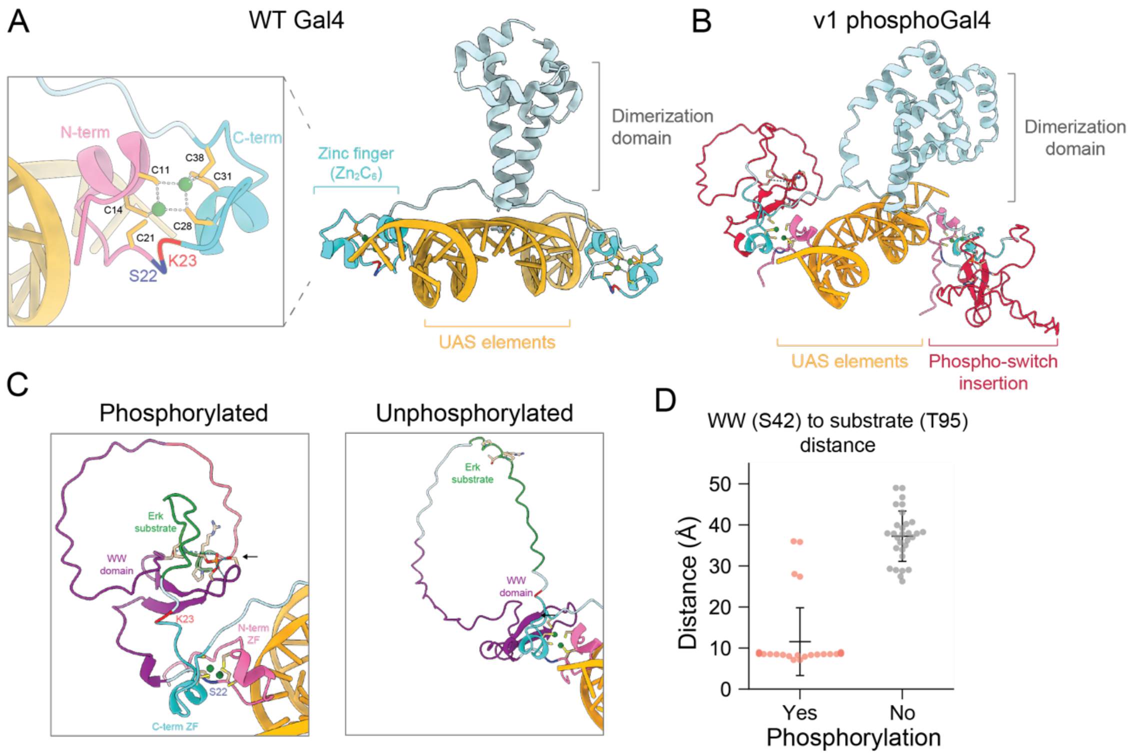


**Supplementary Figure 4. AlphaFold3 predicted structural models of ERK-phosphoGal4v1.** (**A**) Crystal structure of wildtype Gal4 homodimer-DNA complex (PDB: 3COQ). Detailed structure of $Zn_2C_6$ zinc finger of Gal4 shows 6 cysteines (C11, C14, C21, C28, C31, C38) capturing 2 zinc ions (green). Allosteric insertion site between S22 (dark blue) and K23 (red) separates the zinc finger to N-term half (pink) and C-term half (light blue). (**B**) AlphaFold3 predicted models of ERK phospho-switch v1 (dark red) inserted Gal4 in complex with UAS elements with ERK targeted T95 phosphorylation. (**C**) Comparison of phosphorylated and unphosphorylated AlphaFold3 models. Arrow shows contacts between phosphorylated T95 in ERK substrate (green) and the binding interface in the Pin1 WW domain (purple). (**D**) Distances between S42 in the WW binding interface and T95 in the ERK substrate are measured in 15 predicted models (30 data points for each model contains 2 zinc fingers as Gal4 forms a dimer when binds to UAS elements).

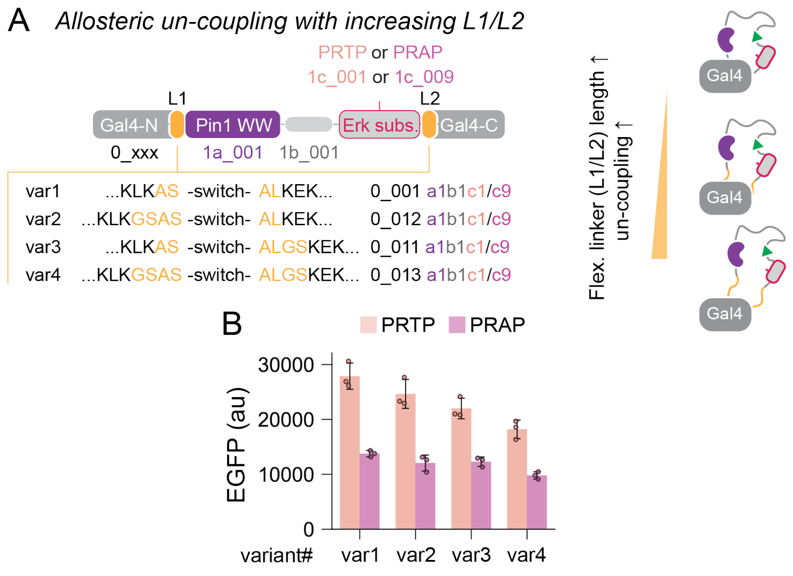


**Supplementary Figure 5. Allosteric un-coupling with ERK-phosphoGal4v1.** (**A**) Generation of ERK-phosphoGal4v1 variants with increasing length of flexible linkers (L1/L2) between ERK phospho-switches and Gal4. Variant 1 is the unmodified ERK-phosphoGal4v1. Variant 2-4 add two-residue GS linkers at N-term side, C-term side or both sides of ERK phospho-switch v1. Increasing length of L1/L2 un-couples the phospho-switch from target protein function. Variants were generated with QCTK Golden Gate assembly. (**B**) GFP expression of ERK phosphoGal4v1 variants 1-4. For each variant, a non-phosphorylatable mutant was tested as control. Cells transiently expressing variants were stimulated with four GM changes in 30 h before flow cytometry measurements. Error bars represent mean± s.e.m (n = 3 replicates).

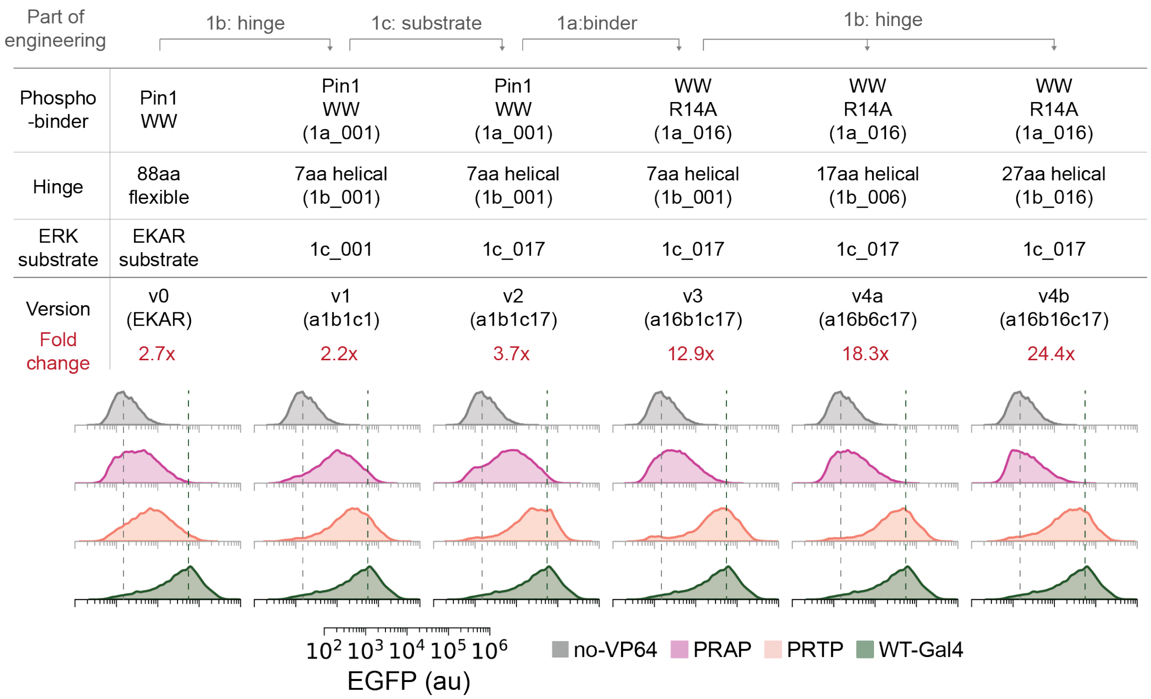


**Supplementary Figure 6. Course of ERK phospho-switch optimization.** Histograms show the performance of ERK-phosphoGal4 (controlled by corresponding versions of ERK phospho-switches) with a WT non-inserted Gal4 (green) and a no-VP64 version of ERK phosphoGal4v3 (gray) as controls. The performance is presented by comparing GFP expressions of the phosphorylatable (PRTP, orange) and non-phosphorylatable (PRAP, pink) phosphoGal4 variants in ERK-ON condition. Histograms represent >10000 cells combined from 3-5 replicates. Cells are gated for IRES_mCherry expression range of ($5\times10^4$-$1.25\times10^5$) for fold change calculation and histograms (**Fig. S1**; See **Methods**). Top charts summarize the course of optimization.

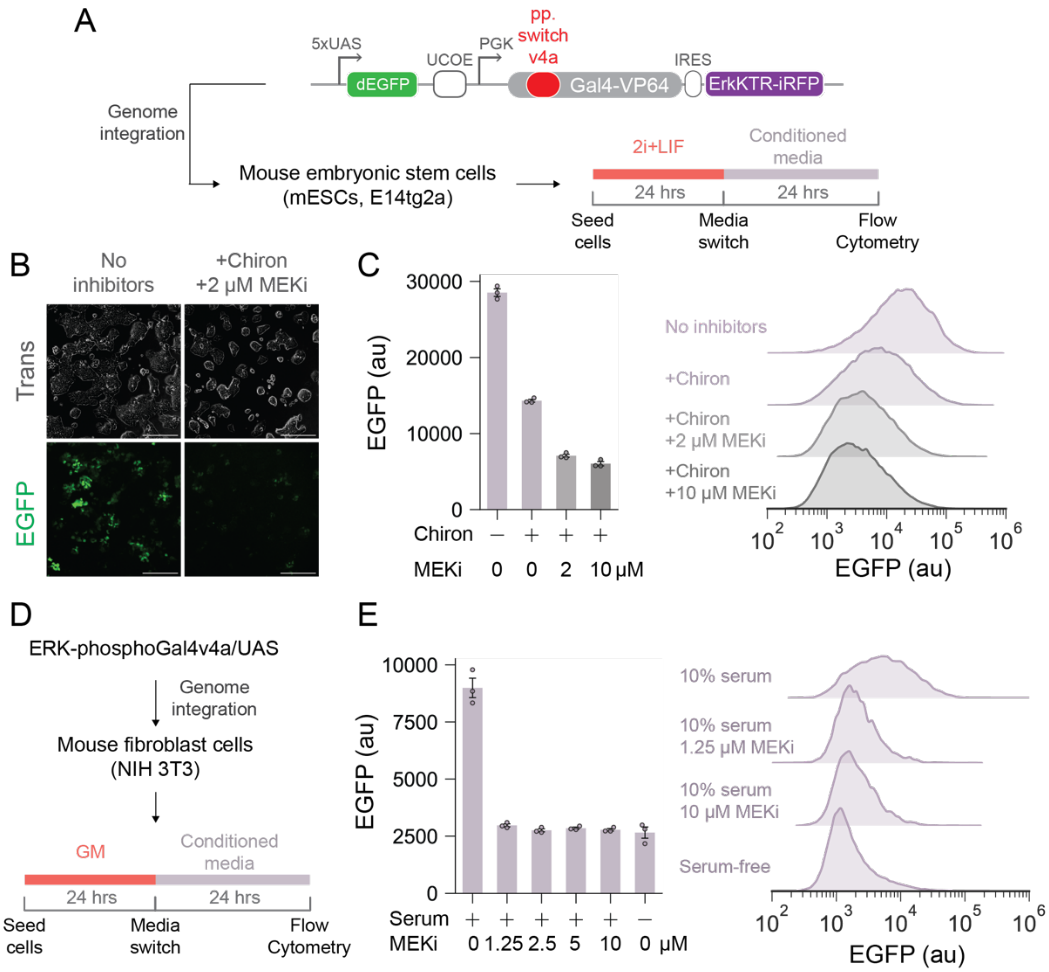


**Supplementary Figure 7. ERK activity biosensing in stem cells and fibroblasts with ERK-phosphoGal4v4a.** (**A**) Schematic of single-construct ERK-phosphoGal4v4a/UAS/ErkKTR system. Mouse embryonic stem cells (mESCs) were engineered to stably express the system. Engineered cells were seeded in 2i+LIF media and switched to the conditioned media for 24 h. (**B**) Representative images of stem cell colony morphology and GFP expression in LIF-only media (No inhibitors) or in 2i+LIF media. Scale bar, 250 μm. (**C**) GFP expression of engineered mESCs in response to drug treatments. Chiron (CHIR-99021), 3 μM. MEKi, PD0325901. Error bars represent mean ± s.e.m (n = 3 replicates). Histograms represent >10000 cells combined from 3 replicates. (**D**) Schematic of NIH 3T3 cells engineered with the single-construct system in **A**. Engineered cells were seeded in growth media (contains 10% FBS) and switched to the conditioned media for 24 h. (**E**) GFP expression of engineered NIH 3T3 cells in response to drug treatments. Serum, 10% FBS. MEKi, PD0325901. Error bars represent mean ± s.e.m (n = 3 replicates). Histograms represent >10000 cells combined from 3 replicates.

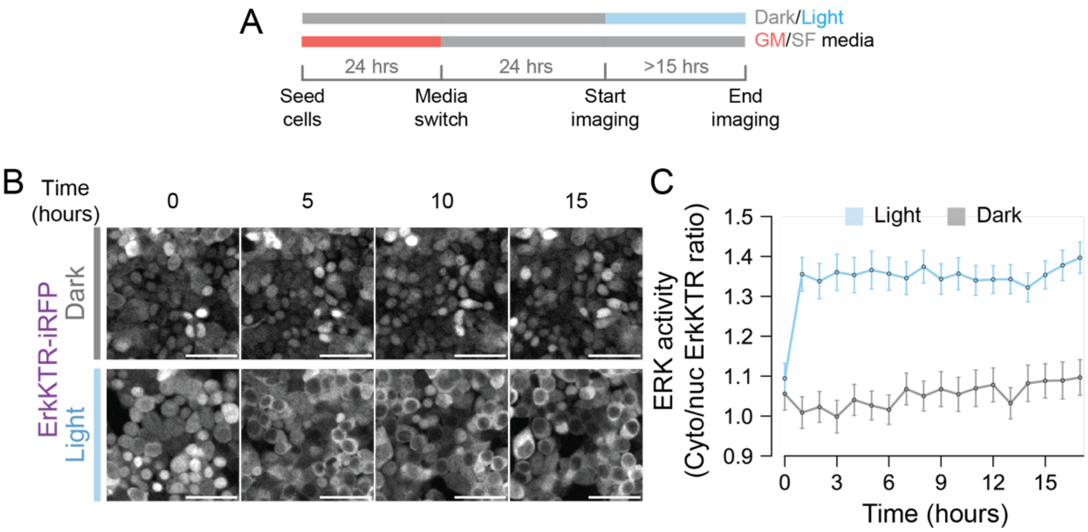


**Supplementary Figure 8. Optogenetic activation of ERK over long time periods.** (**A**) Schematic of live-cell imaging experiments for long-time optogenetic ERK stimulation. Engineered optoSOS/ERK-phosphoGal4v2/UAS HEK293T cells were seeded in growth media (GM) and then switched to serum-free (SF) media for 24 h. Cells were kept in dark before imaging. After starvation, cells were imaged and stimulated with blue light for >15 h. (See **Methods**). (**B**) Representative ErkKTR images at 0, 5, 10 and 15 h in dark or under blue light illumination. Scale bar, 100 µm. (**C**) Quantification of ErkKTR response to long-term optogenetic ERK stimulation. Mean trajectories (solid lines) are shown with error bars representing the 95% confidence interval. Average ~250 cells are quantified at each time points.

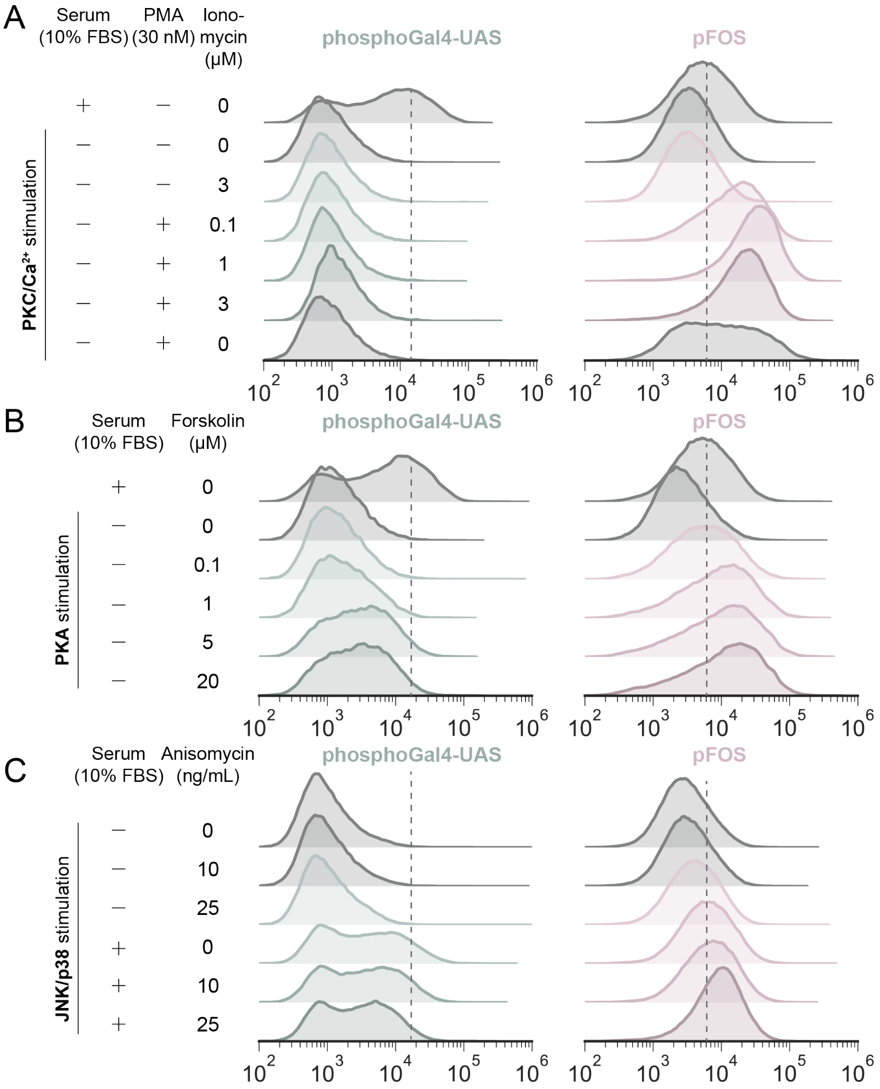


**Supplementary Figure 9. Histograms for ERK-phosphoGal4v4a/UAS and *$P_{FOS}$* promoter comparison.** (**A**) Histograms of GFP expression in response to PKC/$Ca^{2+}$ related activators PMA and ionomycin. (**B**) Histograms of GFP expression in response to the PKA activators Forskolin. (**C**) Histograms of GFP expression in response to the JNK/p38 activator anisomycin.

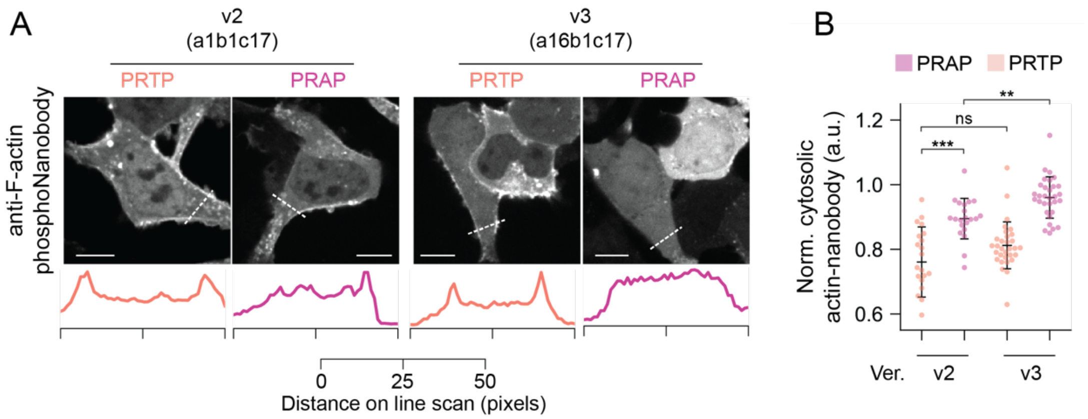


**Supplementary Figure 10. Comparison of F-actin ERK-phosphoNanobody with v2 or v3 ERK phospho-switches.** (**A**) Representative images of ERK-phosphoNanobody localization 7 mins after serum stimulation. For each version, a non-phosphorylatable (PRAP) variant was tested as control. A 50-pixel line scan (11.8 µm) was drawn to visualize the cell periphery localization of tagRFP-tagged ERK-phosphoNanobodies. Scale bar, 10 µm. (**B**) Quantification of ERK-phosphoNanobody cytosolic clearance in response to serum stimulation. Error bars represent mean ± s.d (v2 PRTP, n = 21 cells; v2 PRAP, n = 21 cells; v3 PRTP, n = 31 cells; v3 PRAP, n = 31 cells. Cells are quantified from 2-3 independent experiments). Statistical significance is verified with Mann-Whitney-Wilcoxon tests (two-sided with Bonferroni correction). p-value annotation: ns, $p > 0.05$; *, $0.01 < p <= 0.05$; **, $10^{-3} < p <= 0.01$; ***, $10^{-4} < p <= 10^{-3}$; ****, $p <= 10^{-4}$.

**Supplementary Table 1. List of QCTK1b parts**

| Part name | Part | Cat. | Description (length) | Hinge sequence |
|---|---|---|---|---|
| QCTK1b_001 | 1b | Syn. | EAAAK-repeat helical (7aa) | EAAAKEA |
| QCTK1b_002 | 1a | Nat. | GS-rich flexible (48aa) | SAGGSAGGSAGGSAGGSAGGSGSAGGS AGGSTSAGGSAGGSAGGSGSA |
| QCTK1b_003 | 1b | Nat. | Ja helix of AsLOV2 (26aa) | EHVRDAAEREGVMLIKKTAENIDEAA |
| QCTK1b_004 | 1b | Nat. | tonB (33aa) | EPEPEPEPIPEPPKEAPVVIEKPKPKPKPK PKP |
| QCTK1b_005 | 1b | Syn. | AP-repeat Pro-rich (7aa) | APAPAPA |
| QCTK1b_006 | 1b | Syn. | EAAAK-repeat helical (17aa) | EAAAKEAAAKEAAAKEA |
| QCTK1b_007 | 1b | Nat. | NES (13aa) | DELLKELADLNLD |
| QCTK1b_008 | 1b | Syn. | AP-repeat Pro-rich (17aa) | APAPAPAPAPAPAPAPA |
| QCTK1b_009 | 1b | Nat. | ssrA (8aa) | AANDENYF |
| QCTK1b_010 | 1b | Syn. | GS-rich flexible (17aa) | SAGGSAGGSAGGSAGGS |
| QCTK1b_011 | 1b | Syn. | GS-rich flexible (7aa) | SAGGSAG |
| QCTK1b_012 | 1b | Syn. | AP-repeat Pro-rich (27aa) | APAPAPAPAPAPAPAPAPAPAPAPAPA |
| QCTK1b_013 | 1b | Syn. | EAAAK/GS 50% mix (17aa) | ESAAGSAGAKEAGGKEA |
| QCTK1b_014 | 1b | Syn. | EAAAK/GS 50% mix (7aa) | GSAAKGA |
| QCTK1b_015 | 1b | Syn. | GS/AP 50% mix (7aa) | SPGGPAP |
| QCTK1b_016 | 1b | Syn. | EAAAK-repeat helical (27aa) | EAAAKEAAAKEAAAKEAAAKEAAAKE A |
| QCTK1b_017 | 1b | Syn. | GS-rich flexible (27aa) | SAGGSAGGSAGGSAGGSAGGSGSAGGS |
| QCTK1b_018 | 1b | Syn. | EAAAK/AP 50% mix (17aa) | PAPAKEPPPKPAAAPEP |
| QCTK1b_019 | 1b | Syn. | EAAAK/AP 50% mix (7aa) | EAPAPEP |

| | | | | |
|---|---|---|---|---|
| QCTK1b_020 | 1b | Syn. | EAAAK/AP 50% mix (27aa) | PPAAKPAAAPPAAAKPPPPKPAPAKPP |
| QCTK1b_021 | 1b | Syn. | GS/AP 50% mix (17aa) | SAPGPAGGPPGGPPGPP |
| QCTK1b_022 | 1b | Nat. | HLH fof cI repressor (20aa) | LGLSQESVADKMGMGQSGVG |
| QCTK1b_023 | 1b | Syn. | GS/AP 50% mix (27aa) | PAPGSPGPSAGPPPPGSPPGPGSAPGP |
| QCTK1b_024 | 1b | Nat. | bHLH from human MyoD (52aa) | DRRKAATMRERRRLSKVNEAFETLKRC TSSNPNQRLPKVEILRNAIRYIEGL |
| QCTK1b_025 | 1b | Syn. | EAAAK/GS 50% mix (27aa) | GAGASEASAKEGGAGSGAAKEAAAKE A |
| QCTK1b_026 | 1b | Nat. | Upper hinge of human IgG (12aa) | EPKSCDKTHTCP |

**Supplementary Table 2. List of QCTK1a/c parts**

| Part name | Part | Description | Protein sequence (if applicable*) |
|---|---|---|---|
| QCTK0_001 | 0 | Used to generate ERK-phophoGal4v0-1 | / |
| QCTK0_001_v2 | 0 | Used to generate ERK-phophoGal4v2-4a/b | / |
| QCTK0_011 | 0 | Used to generate L1/L2 phosphoGal4 variants in **Fig. S5** | / |
| QCTK0_012 | 0 | Used to generate L1/L2 phosphoGal4 variants in **Fig. S5** | / |
| QCTK0_013 | 0 | Used to generate L1/L2 phosphoGal4 variants in **Fig. S5** | / |
| QCTK1a_001 | 1a | Pin1 WW domain | MADEEKLPPGWEKRMSRSSGRVYYFNHITNASQWERPSGNSSSGGKNGQGEPAR |
| QCTK1a_014 | 1a | WW domain N-term MADE-deletion | EKLPPGWEKRMSRSSGRVYYFNHITNASQWERPSGNSSSGGKNGQGEPAR |
| QCTK1a_015 | 1a | WW domain N-term MADEEKLPP-deletion | GWEKRMSRSSGRVYYFNHITNASQWERPSGNSSSGGKNGQGEPAR |
| QCTK1a_016 | 1a | WW R14A | MADEEKLPPGWEKAMSRSSGRVYYFNHITNASQWERPSGNSSSGGKNGQGEPAR |
| QCTK1a_017 | 1a | WW F25L | MADEEKLPPGWEKRMSRSSGRVYYLNHITNASQWERPSGNSSSGGKNGQGEPAR |
| QCTK1c_001 | 1c | ERK substrate from EKAR | LPDVPRTPVDKAKLSFQFP |
| QCTK1c_004 | 1c | JNK substrate | DSVKTPEDEGNPLLEQLEKK |
| QCTK1c_008 | 1c | Used to generate Subs. 2 in **Fig. 3A**** | LPDVPRTPVDKAKLSFQFP |
| QCTK1c_009 | 1c | T-to-A mutated ERK substrate from EKAR | LPDVPRAPVDKAKLSFQFP |
| QCTK1c_014 | 1c | Subs. 5 in **Fig. 3A** | LAAVARAAVAKAKLAFQFP |
| QCTK1c_016 | 1c | Subs. 4 in **Fig. 3A** | LPDVPRAPVDKAKLAFQFP |
| QCTK1c_017 | 1c | ERK substrate for ERK phospho-switch v2 | LAAVPRTPVAKAKLAFQFP |
| QCTK1c_018 | 1c | T-to-A mutated ERK substrate for ERK phospho-switch v2 | LAAVPRAPVAKAKLAFQFP |

* For Level 1a/1c parts, protein sequences are provided.
** 1c_008 and 1c_001 share the same substrate sequence. 1c_008 part has modified BsaI overhangs and after assembly, the result phosphoGal4 do not contain Ser in the linkers between the phospho-switch and Gal4 (as showed in **Fig. 3A**).

**Supplementary Table 3. List of important plasmids used in this study**

| Plasmid name | Figure | Assembly | Description |
|---|---|---|---|
| 0_001_a1b10c1 | 2 | Golden Gate | PB_EF1a_ERK_phosphoGal4(0_001_a1b10c1)_IRES_mCherry |
| 0_001_a1b11c1 | 2 | Golden Gate | PB_EF1a_ERK_phosphoGal4(0_001_a1b11c1)_IRES_mCherry |
| 0_001_a1b12c1 | 2 | Golden Gate | PB_EF1a_ERK_phosphoGal4(0_001_a1b12c1)_IRES_mCherry |
| 0_001_a1b13c1 | 2 | Golden Gate | PB_EF1a_ERK_phosphoGal4(0_001_a1b13c1)_IRES_mCherry |
| 0_001_a1b14c1 | 2 | Golden Gate | PB_EF1a_ERK_phosphoGal4(0_001_a1b14c1)_IRES_mCherry |
| 0_001_a1b15c1 | 2 | Golden Gate | PB_EF1a_ERK_phosphoGal4(0_001_a1b15c1)_IRES_mCherry |
| 0_001_a1b16c1 | 2 | Golden Gate | PB_EF1a_ERK_phosphoGal4(0_001_a1b16c1)_IRES_mCherry |
| 0_001_a1b17c1 | 2 | Golden Gate | PB_EF1a_ERK_phosphoGal4(0_001_a1b17c1)_IRES_mCherry |
| 0_001_a1b18c1 | 2 | Golden Gate | PB_EF1a_ERK_phosphoGal4(0_001_a1b18c1)_IRES_mCherry |
| 0_001_a1b19c1 | 2 | Golden Gate | PB_EF1a_ERK_phosphoGal4(0_001_a1b19c1)_IRES_mCherry |
| 0_001_a1b1c1 | 2 | Golden Gate | PB_EF1a_ERK_phosphoGal4(0_001_a1b1c1)_IRES_mCherry |
| 0_001_a1b1c9 | S5 | Golden Gate | PB_EF1a_ERK_phosphoGal4(0_001_a1b1c9)_IRES_mCherry |
| 0_001_a1b20c1 | 2 | Golden Gate | PB_EF1a_ERK_phosphoGal4(0_001_a1b20c1)_IRES_mCherry |
| 0_001_a1b21c1 | 2 | Golden Gate | PB_EF1a_ERK_phosphoGal4(0_001_a1b21c1)_IRES_mCherry |
| 0_001_a1b22c1 | 2 | Golden Gate | PB_EF1a_ERK_phosphoGal4(0_001_a1b22c1)_IRES_mCherry |
| 0_001_a1b23c1 | 2 | Golden Gate | PB_EF1a_ERK_phosphoGal4(0_001_a1b23c1)_IRES_mCherry |
| 0_001_a1b24c1 | 2 | Golden Gate | PB_EF1a_ERK_phosphoGal4(0_001_a1b24c1)_IRES_mCherry |
| 0_001_a1b25c1 | 2 | Golden Gate | PB_EF1a_ERK_phosphoGal4(0_001_a1b25c1)_IRES_mCherry |
| 0_001_a1b26c1 | 2 | Golden Gate | PB_EF1a_ERK_phosphoGal4(0_001_a1b26c1)_IRES_mCherry |
| 0_001_a1b2c1 | 2 | Golden Gate | PB_EF1a_ERK_phosphoGal4(0_001_a1b2c1)_IRES_mCherry |
| 0_001_a1b3c1 | 2 | Golden Gate | PB_EF1a_ERK_phosphoGal4(0_001_a1b3c1)_IRES_mCherry |
| 0_001_a1b4c1 | 2 | Golden Gate | PB_EF1a_ERK_phosphoGal4(0_001_a1b4c1)_IRES_mCherry |
| 0_001_a1b5c1 | 2 | Golden Gate | PB_EF1a_ERK_phosphoGal4(0_001_a1b5c1)_IRES_mCherry |
| 0_001_a1b6c1 | 2 | Golden Gate | PB_EF1a_ERK_phosphoGal4(0_001_a1b6c1)_IRES_mCherry |
| 0_001_a1b7c1 | 2 | Golden Gate | PB_EF1a_ERK_phosphoGal4(0_001_a1b7c1)_IRES_mCherry |
| 0_001_a1b8c1 | 2 | Golden Gate | PB_EF1a_ERK_phosphoGal4(0_001_a1b8c1)_IRES_mCherry |
| 0_001_a1b9c1 | 2 | Golden Gate | PB_EF1a_ERK_phosphoGal4(0_001_a1b9c1)_IRES_mCherry |

| | | | |
|---|---|---|---|
| 0_001_v2_a16b16c17 | 3F | Golden Gate | PB_EF1a_ERK_phosphoGal4(0_001_v2_a16b16c17)_IRES_mCherry |
| 0_001_v2_a16b16c18 | 3F | Golden Gate | PB_EF1a_ERK_phosphoGal4(0_001_v2_a16b16c18)_IRES_mCherry |
| 0_001_v2_a16b1c17 | 3E | Golden Gate | PB_EF1a_ERK_phosphoGal4(0_001_v2_a16b1c17)_IRES_mCherry |
| 0_001_v2_a16b1c18 | 3E | Golden Gate | PB_EF1a_ERK_phosphoGal4(0_001_v2_a16b1c18)_IRES_mCherry |
| 0_001_v2_a16b6c17 | 3F | Golden Gate | PB_EF1a_ERK_phosphoGal4(0_001_v2_a16b6c17)_IRES_mCherry |
| 0_001_v2_a16b6c18 | 3F | Golden Gate | PB_EF1a_ERK_phosphoGal4(0_001_v2_a16b6c18)_IRES_mCherry |
| 0_001_v2_a17b1c17 | 3E | Golden Gate | PB_EF1a_ERK_phosphoGal4(0_001_v2_a17b1c17)_IRES_mCherry |
| 0_001_v2_a17b1c18 | 3E | Golden Gate | PB_EF1a_ERK_phosphoGal4(0_001_v2_a17b1c18)_IRES_mCherry |
| 0_001_v2_a1b1c14 | 3A-B | Golden Gate | PB_EF1a_ERK_phosphoGal4(0_001_v2_a1b1c14)_IRES_mCherry |
| 0_001_v2_a1b1c16 | 3A-B | Golden Gate | PB_EF1a_ERK_phosphoGal4(0_001_v2_a1b1c16)_IRES_mCherry |
| 0_001_v2_a1b1c17 | 3C | Golden Gate | PB_EF1a_ERK_phosphoGal4(0_001_v2_a1b1c17)_IRES_mCherry |
| 0_001_v2_a1b1c18 | 3C | Golden Gate | PB_EF1a_ERK_phosphoGal4(0_001_v2_a1b1c18)_IRES_mCherry |
| 0_001_v2_a1b1c4 | 6C | Golden Gate | PB_EF1a_JNK_phosphoGal4(0_001_v2_a1b1c4)_IRES_mCherry |
| 0_001_v2_a1b1c8 | 3A-B | Golden Gate | PB_EF1a_ERK_phosphoGal4(0_001_v2_a1b1c8)_IRES_mCherry |
| 0_011_a1b1c1 | S5 | Golden Gate | PB_EF1a_ERK_phosphoGal4(0_011_a1b1c1)_IRES_mCherry |
| 0_011_a1b1c9 | S5 | Golden Gate | PB_EF1a_ERK_phosphoGal4(0_011_a1b1c9)_IRES_mCherry |
| 0_012_a1b1c1 | S5 | Golden Gate | PB_EF1a_ERK_phosphoGal4(0_012_a1b1c1)_IRES_mCherry |
| 0_012_a1b1c9 | S5 | Golden Gate | PB_EF1a_ERK_phosphoGal4(0_012_a1b1c9)_IRES_mCherry |
| 0_013_a1b1c1 | S5 | Golden Gate | PB_EF1a_ERK_phosphoGal4(0_013_a1b1c1)_IRES_mCherry |
| 0_013_a1b1c9 | S5 | Golden Gate | PB_EF1a_ERK_phosphoGal4(0_013_a1b1c9)_IRES_mCherry |
| pQC061 | 1 | In-Fusion | PB_EF1a_Gal4_VP64_EKAR_88aa_IRES_mCherry |
| pQC086 | 1 | In-Fusion | PB_EF1a_Gal4_VP64_EKAR_88aa_TA_IRES_mCherry |
| pQC131 | 4 | In-Fusion | PB_UAS_dEGFP_PGK_iRFP_ErkKTR |
| pQC136 | 4 | In-Fusion | PB_EF1a_tagBFP_SSPB_SOScat_P2A_iLID_CAAX |
| pQC137 | 5 | In-Fusion | PB_pFOS_dEGFP_PGK_iRFP_ErkKTR |
| pQC188 | 6C | In-Fusion | PB_EF1a_Gal4_a1b1c4_TA_IRES_mCherry |
| pQC234 | 3A-B | In-Fusion | PB_EF1a_Gal4(0_001v2_a1b1c8)_with_SA_mutation (as the Subs. 3 variant) |
| pQC235 | 6H-J | In-Fusion | pHR_SFFV_ActinNanobody_switch(a16b1c17)_tagRFP |

| | | | |
|---|---|---|---|
| pQC236 | 6H-J | In-Fusion | pHR_SFFV_ActinNanobody_switch(a16b1c18)_tagRFP |
| pQC238 | 3G | In-Fusion | PB_EF1a_Gal4(a16b1c17)_no_VP64_IRES_mCherry_ctrl |
| pQC256 | 6C | In-Fusion | PB_EF1a_Gal4_a16b1c4_IRES_mCherry |
| pQC257 | 6C | In-Fusion | PB_EF1a_Gal4_a16b1c4_TA_IRES_mCherry |
| pQC274 | S8 | In-Fusion | PB-UAS-dGFP-UCOE-PGK-Gal4(0_001v2_a16b6c17)-VP64-IRES-ErkKTR-iRFP |
| pQC060 | 1 | In-Fusion | PB_EF1a_Gal4_VP64_IRES_mCherry |
| pQC171 | S10 | In-Fusion | pHR_SFFV_ActinNanobody_switch(a1b1c17)_tagRFP |
| pQC172 | S10 | In-Fusion | pHR_SFFV_ActinNanobody_switch(a1b1c18)_tagRFP |

**Supplementary Table 4. List of important protein sequences in this study**

| Name | Category | Protein sequence* |
|---|---|---|
| ERK phospho-switch v0 (EKAR) | Phospho-switch | MADEEKLPPGWEKRMSRSSGRVYYFNHITNASQWER<br>PSGNSSSGGKNGQGEPARGTSAGGSAGGSAGGSAGGS<br>AGGSGSAGGSAGGSTSAGGSAGGSAGGSAGGSAGGS<br>GSAGGSAGGSTSAGGSAGGSAGGSAGGSAGGSGSAG<br>SPDVPRTPVDKAKLSFQFP |
| ERK phospho-switch v1 (a1b1c1) | Phospho-switch | MADEEKLPPGWEKRMSRSSGRVYYFNHITNASQWER<br>PSGNSSSGGKNGQGEPARGTEAAAKEAGSPDVPRTPV<br>DKAKLSFQFP |
| ERK phospho-switch v2 (a1b1c17) | Phospho-switch | MADEEKLPPGWEKRMSRSSGRVYYFNHITNASQWER<br>PSGNSSSGGKNGQGEPARGTEAAAKEAGLAAVPRTPV<br>AKAKLAFQFP |
| ERK phospho-switch v3 (a16b1c17) | Phospho-switch | MADEEKLPPGWEKAMSRSSGRVYYFNHITNASQWER<br>PSGNSSSGGKNGQGEPARGTEAAAKEAGLAAVPRTPV<br>AKAKLAFQFP |
| ERK phospho-switch v4a (a16b6c17) | Phospho-switch | MADEEKLPPGWEKAMSRSSGRVYYFNHITNASQWER<br>PSGNSSSGGKNGQGEPARGTEAAAKEAAAKEAAAKE<br>AGLAAVPRTPVAKAKLAFQFP |
| ERK phospho-switch v4b (a16b16c17) | Phospho-switch | MADEEKLPPGWEKAMSRSSGRVYYFNHITNASQWER<br>PSGNSSSGGKNGQGEPARGTEAAAKEAAAKEAAAKE<br>AAAKEAAAKEAGLAAVPRTPVAKAKLAFQFP |
| JNK phospho-switch v3 (a16b1c4) | Phospho-switch | MADEEKLPPGWEKAMSRSSGRVYYFNHITNASQWER<br>PSGNSSSGGKNGQGEPARGTEAAAKEAGLDSVKTPED<br>EGNPLLEQLEKK |
| ERK-phosphoGal4v4a | Phospho-protein | MKLLSSIEQACDICRLKKLKCSASMADEEKLPPGWEK<br>AMSRSSGRVYYFNHITNASQWERPSGNSSSGGKNGQ<br>GEPARGTEAAAKEAAAKEAAAKEAGLAAVPRTPVAK<br>AKLAFQFPALKEKPKCAKCLKNNWECRYSPKTKRSPL<br>TRAHLTEVESRLERLEQLFLLIFPREDLDMILKMDSLQ<br>DIKALLTGLFVQDNVNKDAVTDRLASVETDMPLTLR<br>QHRISATSSSEESSNKGQRQLTVSAAAGGSGGSGGSD<br>ALDDFDLDMLGSDALDDFDLDMLGSDALDDFDLDM<br>LGSDALDDFDLDMLGS |
| ERK-phosphoGal4v4b | Phospho-protein | MKLLSSIEQACDICRLKKLKCSASMADEEKLPPGWEK<br>AMSRSSGRVYYFNHITNASQWERPSGNSSSGGKNGQ<br>GEPARGTEAAAKEAAAKEAAAKEAAAKEAAAKEAG<br>LAAVPRTPVAKAKLAFQFPALKEKPKCAKCLKNNWE<br>CRYSPKTKRSPLTRAHLTEVESRLERLEQLFLLIFPRED<br>LDMILKMDSLQDIKALLTGLFVQDNVNKDAVTDRLA<br>SVETDMPLTLRQHRISATSSSEESSNKGQRQLTVSAAA<br>GGSGGSGGSDALDDFDLDMLGSDALDDFDLDMLGSD<br>ALDDFDLDMLGSDALDDFDLDMLGS |
| JNK-phosphoGal4v3 | Phospho-protein | MKLLSSIEQACDICRLKKLKCSASMADEEKLPPGWEK<br>AMSRSSGRVYYFNHITNASQWERPSGNSSSGGKNGQ<br>GEPARGTEAAAKEAGLDSVKTPEDEGNPLLEQLEKKA<br>LKEKPKCAKCLKNNWECRYSPKTKRSPLTRAHLTEVE<br>SRLERLEQLFLLIFPREDLDMILKMDSLQDIKALLTGLF<br>VQDNVNKDAVTDRLASVETDMPLTLRQHRISATSSSE<br>ESSNKGQRQLTVSAAAGGSGGSGGSDALDDFDLDML<br>GSDALDDFDLDMLGSDALDDFDLDMLGSDALDDFDL<br>DMLGS |

| | | |
|---|---|---|
| F-actin ERK phosphoNanobodyv3 | Phospho-protein | MAQVQLVESGGGLTQAGGSLRLSCATSGLIFSAFGMGWFRQAPGKEREFVGGINWRGSTNYGDMADEEKLPPGWEKAMSRSSGRVYYFNHITNASQWERPSGNSSSGGKNGQGEPARGTEAAAKEAGLAAVPRTPVAKAKLAFQFPGRFTISRDNAKNTVYLQMNNLKPEDTAVYYCAARMVHKTEYDYWGEGTQVTVSSRSLGGGGSGGGGSGGGGSGGGGSMVSKGEELIKENMHMKLYMEGTVNNHHFKCTSEGEGKPYEGTQTMRIKVVEGGPLPFAFDILATSFMYGSRTFINHTQGIPDFFKQSFPEGFTWERVTTYEDGGVLTATQDTSLQDGCLIYNVKIRGVNFPSNGPVMQKKTLGWEANTEMLYPADGGLEGRSDMALKLVGGGHLICNFKTTYRSKKPAKNLKMPGVYYVDHRLERIKEADKETYVEQHEVAVARYCDLPSKLGHKLN |

* Color scheme for the protein sequence annotation. Purple: Phospho-binding domain; Gray: Hinge; Yellow: Kinase substrate; Light blue: Gal4; Orange: VP64; Dark blue: Actin Nanobody; Dark red: TagRFP
